# Spatial tissue profiling by imaging-free molecular tomography

**DOI:** 10.1101/2020.08.04.235655

**Authors:** Halima Hannah Schede, Christian G. Schneider, Johanna Stergiadou, Lars E. Borm, Anurag Ranjak, Tracy M. Yamawaki, Fabrice P.A. David, Peter Lönnerberg, Gilles Laurent, Maria Antonietta Tosches, Simone Codeluppi, Gioele La Manno

## Abstract

Genomics techniques are currently being adapted to provide spatially resolved omics profiling. However, the adaptation of each new method typically requires the setup of specific detection strategies or specialized instrumentation. A generic approach to spatially resolve different types of high throughput data is missing. Here, we describe an imaging-free framework to localize high throughput readouts within a tissue by combining compressive sampling and image reconstruction. We implemented this framework to transform a low-input RNA sequencing protocol into an imaging-free spatial transcriptomics technique (STRP-seq) and validated this method with a transcriptome profiling of the murine brain. To verify the broad applicability of STRP-seq, we applied the technique on the brain of the Australian bearded dragon *Pogona vitticeps*. Our results reveal the molecular anatomy of the telencephalon of this lizard, providing evidence for a marked regionalization of the reptilian pallium and subpallium. Overall, the proposed framework constitutes a new approach that allows upgrading in a generic fashion conventional genomic assays to spatially resolved techniques.

## INTRODUCTION

Spatially resolved molecular atlases constitute fundamental resources to the scientific community [1]. In situ hybridization (ISH) compendia such as the Allen Brain Atlas or Genepaint are extensively used as references to guide the mapping of newly discovered cell types [2]–[6]. Recently, the development of new technologies that can simultaneously map the expression of multiple genes on the same tissue sample have accelerated the generation of new atlases. These techniques are based on either quantitative in-situ hybridization or on the sequencing of RNA molecules captured on barcoded supports [7]–[17]. However, in-situ hybridization techniques require a preselection of target genes, and sequencing methods suffer from resolution limits and suboptimal capture efficiency.

An alternative approach for spatially resolving the distribution of gene expression patterns relies on sampling a tissue regularly on a grid by laser capture microdissection (LCM). After LCM sampling, a next-generation sequencing (NGS) library can be prepared using a low-input protocol, and the gene expression pattern can be quantified [18]. Overall, combining microdissection-based approaches with single-cell RNA sequencing has proved successful to delineate relationships between cells populating a niche and their potency [19]. However, the sensitivity of LCM-based assays is limited by laser-induced degradation of nucleic-acids and the technical complexity of handling and preparing libraries for multiple microscopic samples is considerable.

Tomo-seq is a more accessible and imaging-free spatial sequencing method in which a tissue sample is cryosectioned into thin slices that are used to prepare NGS libraries [20], [21]. Using this method, it is possible to generate a one-dimensional transcriptomic pattern that allows the evaluation of gene expression variation over a spatial axis. The authors also demonstrated the reconstruction of three-dimensional gene expression patterns by using the iterative proportional fitting (IPF) algorithm to combine three orthogonal one-dimensional profiles generated using multiple genetically equivalent samples [20]. However, IPF cannot solve this reconstruction problem for complex anatomical configurations and lacks robustness to noise. Moreover, an entire specimen is required to obtain each of the 1D profiles, making 3D reconstructions impossible for many applications where multiple genetically identical specimens are not available [22]–[24].

Even though the current atlassing efforts are focused on mapping gene expression patterns, the possibility to create atlases probing other modalities will be beneficial for the entire scientific community [25], [26]. However, other NGS-based technologies, including those to measure chromatin accessibility, histone modifications, non-coding or nascent RNAs lack a spatial counterpart. Thus, the development of a generalized approach to convert every existing NGS protocol into a spatially resolved technique, without modifying the detection strategy, would accelerate the transition to a spatial-omics paradigm. Motivated by this, we developed a strategy to perform such an adaptation. Here, we present a framework that brings this transition to life by using a compressed sensing tissue sampling strategy based on multi-angle-sectioning and an associated algorithm that enables the reconstruction of complex spatial patterns (see Online Methods). In this study, we demonstrate that our spatial-omics approach allows the reconstruction of spatial expression patterns on a transcriptome-wide scale by benchmarking it against the mouse Allen Institute ISH Mouse Brain atlas [1]. Furthermore, we apply this framework to study the brain of the lizard non-model organism *Pogona vitticeps*. We reveal quantitative aspects of the molecular anatomy of the pallium and comparative relationships between the reptilian and the mammalian brain.

## RESULTS

### The spatial distribution of a signal can be determined by compressed sampling and reconstruction

The objective of the proposed framework is to determine the abundance of a target signal at a given tissue location through the use of a compressed sensing approach [27]. The approach consists in measuring several locations of the tissue collectively so as to simplify tissue sampling while allowing the subsequent reconstruction of the signal at every single point (Fig. 1).

**Figure 1.**
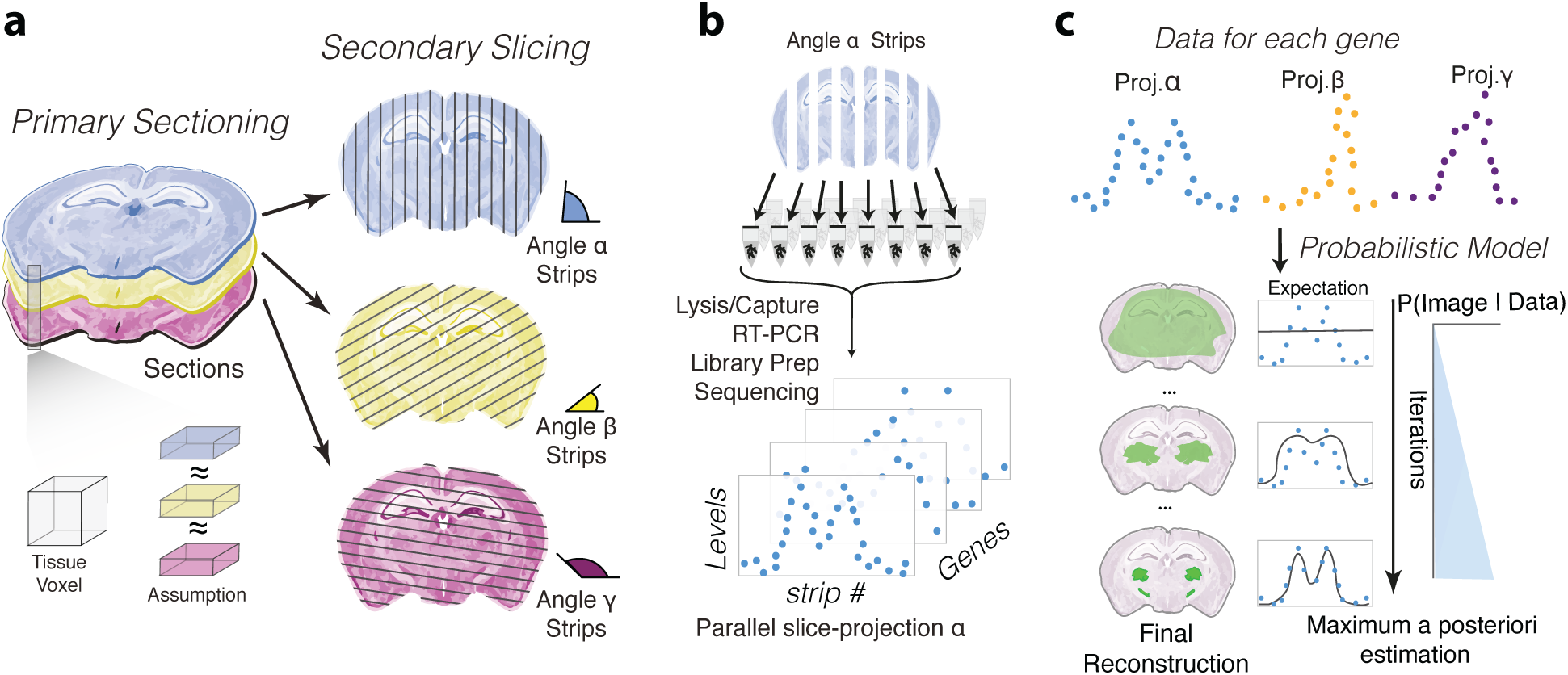
Sampling and reconstruction approach to resolve the spatial localization of genomics data. **a**, Schematic representation of the tissue processing steps. The tissue is first cut along the axis of interest to obtain consecutive 14 µm *primary sections*. The spatial pattern of the signal is assumed constant between primary sections. Each section is then sliced at a different angle to generate tissue strips (*secondary sections*). Only three angles are shown for simplicity, but more are typically used. **b**, Library preparation and generation of raw data. Cells of tissue strips are lysed and mRNA is captured. Reverse transcription is followed by PCR, tagmentation and sequencing. For each primary section, the gene expression within each tissue strip can be concatenated as a parallel-slice projection vector. **c**, Reconstruction of gene expression patterns performed by the *Tomographer* algorithm. Gene expression count values for the strips are modeled as samples from a negative binomial distribution. The expected value is the sum of the intensity of the image to reconstruct. The spatial localization of the gene of interest is obtained by maximum a posteriori estimation, considering probabilistic priors on the distribution of pixel intensities.

To this aim, we designed a user-friendly tissue sampling scheme based on cryosectioning (Supplementary Fig. 1). Initially, the tissue is cut into consecutive thin slices, which we refer to as “primary sections”. Subsequently, primary sections are further sliced along an orthogonal plane at pre-defined orientations resulting in tissue strips (“secondary sections”) (Fig. 1a, see Online Methods). Here, we reason that primary sections are thin enough so that the difference in the signal distribution between them is negligible in relation to the target resolution (i.e. we chose 14µm for the experiments below). Under this assumption, different sets of secondary sections obtained at different orientations can be considered as a resampling of the same distribution.

It is helpful to think of the set of measurements obtained from one series of secondary sections as an analogue of a parallel-beam sum projection in ray-based tomography [28], [29]. Henceforth we refer to this set of measurements for each particular angle as a “parallel-slice projection”. However, in contrast to classical computer tomography, we developed a probabilistic algorithm that requires only a few parallel-slice projections to reconstruct the spatial pattern that generated the data. This is achieved by modeling the noise, setting priors on the degree of smoothness and sparsity of the solution, and then finding the maximum a posteriori estimate (Fig. 1c; see Section 2 of the Online Methods).

We implemented this framework for the quantification of gene expression and developed Spatial Transcriptomics by Reoriented Projections and sequencing (STRP-seq), a method that combines the sampling strategy presented above with a customized low-input RNA-seq protocol based on STRT-seq chemistry (Fig. 1b, see Section 1 of Online Methods) [30]. In this technique, we obtain a parallel-slice projection for each gene by quantifying the reads that map to a transcript in each of the secondary sections (Fig. 1b,c).

### *Tomographer*’s maximum a posteriori estimates provide accurate reconstructions of complex spatial patterns

To demonstrate the potential of the proposed framework for studying non-trivial spatial patterns, we carried out in-silico simulation experiments using the Adult Mouse Allen Brain ISH Atlas profiles as ground truth (see Online Methods Section 3). We compared ground truth ISH profiles to the reconstructions obtained from simulation experiments using both our newly developed algorithm dubbed *Tomographer* and the IPT-based reconstruction proposed for Tomo-seq [20]. Since a 3D reconstruction problem is more strongly underdetermined (i.e. significantly more unknowns than equations) than the one we propose to solve, we include a simple 2D adaptation of Tomo-seq that constitutes a fairer comparison (See Online Methods Section 3.2).

To assess the information that can be recovered by each of the sampling methods, we determined the reconstruction accuracies of genes that are characterized by different sparsity and distribution patterns (Fig. 2a). We calculated the Pearson’s correlation coefficient, the total absolute difference among pixels, and the relative error between the reconstructions and the ground truth. Evaluating these metrics we concluded that our method outperforms Tomo-seq for the reconstruction of the spatial pattern of a given gene (Fig. 2b).

**Figure 2.**
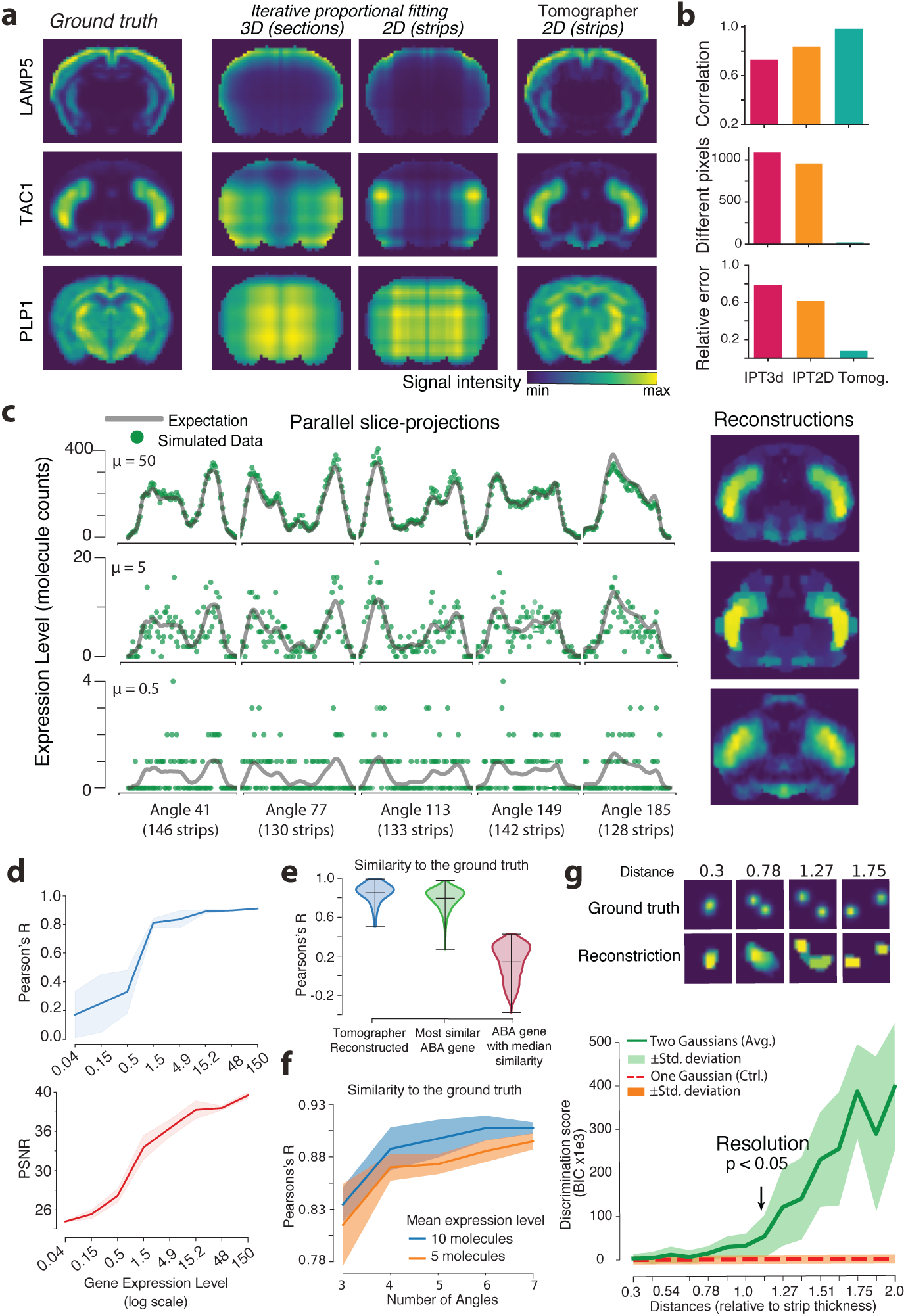
Reconstruction performance of *Tomographer* on simulated data. **a**, Comparison of the novel sampling and reconstruction algorithm (*Tomographer*) with existing methods: Iterative proportional fitting 2D and 3D (Tomo-seq). Starting from gene expression patterns extracted from the Allen Brain Atlas, the simulated projections are computed and the image is then reconstructed (noiseless case). **b**, Reconstruction accuracy for TAC1 evaluated using different metrics between the images: Pearson’s correlation coefficient, relative error and the number of pixels that differ more than the range between the maximum and minimum pixel intensity values are calculated with respect to the ground truth. **c**, Resilience to low signal to noise ratio of the *Tomographer* algorithm. Noisy data are simulated by projecting the expression and adding Poisson noise. **d**, Reconstruction accuracy at different expression levels for a given gene. The relative noise increases as the average expression level decays, causing a decrease in reconstruction accuracy for lowly expressed genes. Pearson correlation coefficient (blue) and Peak signal-to-noise ratio (PSNR) (red) shown for five realizations with additive Poisson noise at various expression levels. The shaded area is the standard deviation over 10 realizations of the noisy projections. **e**, Reconstruction accuracy across a sample of 100 genes of the Allen Brain Atlas with noise simulated as in (c). Reconstructions were compared to the ground truth using Pearson’s correlation coefficient. Violin plots show the distribution of similarity across all the genes. Reference violins show the similarity of each ground truth gene with the most similar gene (green) and the gene median similarity (across the entire Allen Brain Atlas). **f**, Effect of the number of projections (i.e. angles) on the reconstruction accuracy. Reconstructions are scored using Pearson’s correlation coefficient. Standard deviation is computed over different realizations of the noise for average expression levels of 5 and 10. **g**, Assessment of two-points discriminative power and resolution. The two-point effective resolution was defined as the distance at which the two-Gaussian model outperforms the single-Gaussian model in explaining the reconstructed images, the Bayesian Information Criterion (BIC) was used as a metric. Resolution was determined to be 1.15 times the strip thickness (n=16, p-value < 2 × 10^-5). Standard deviation calculated among random realization of the location of the points, fixing the distance.

We developed a probabilistic signal reconstruction strategy taking into account that we expect read counts to be distributed as a Negative Binomial (See Online Methods Section 2; [31], [32]). In this way, we aim at buffering reconstruction errors that may arise because of the discrete nature of the data and the heteroscedasticity within each parallel-slice span. Nevertheless, we expect the maximum achievable accuracy to remain dependent on average expression levels. To test the extent of this dependence, we corrupted simulated parallel-slice projections drawing realizations from a Negative Binomial distribution after rescaling the average read count to different levels (Fig. 2c-e and Supplementary Fig. 2). Analyzing the reconstructions, we found that greater expression levels correlate with the quality of the output and that a minimum average expression level of 1 molecule per strip is required to obtain reliable reconstructions. Notably, the reconstruction accuracy we obtained here was significantly more robust to what we achieved by a naive leastsquares approach (Supplementary Fig. 2a-d)

To gauge the ability of the technique to generalize and resolve other spatial distributions, we simulated reconstructions for 100 randomly selected genes retrieved from the Allen Brain Atlas. The resulting reconstructions were on average more similar to the ground truth pattern of a gene than to the next most similar gene available in the Allen Brain Atlas (Fig. 2e).

We then analyzed the effect of using different numbers of secondary slicing angles on the reconstruction accuracy in order to optimize the reconstruction while maintaining the feasibility of the protocol from an experimental viewpoint. We demonstrate that four cutting angles provide results that are a fair compromise between the reconstruction quality and sample processing effort and cost (Fig. 2f). However, we decided to use five cutting angles in the implementation of the protocol to increase resiliency to other technical sources of error (e.g. cutting errors or lost strips).

Finally, we evaluated the two-point discriminative resolution of the technique in noisy conditions as a function of the thickness of the strip generated by secondary slicing. By performing Monte Carlo simulations we demonstrate that the approach is capable of discriminating two distinct points provided that their distance is at least 1.15 times the secondary section width (Fig. 2g; 80 μm in this case; See Online Methods section 3.6).

### STRP-seq accurately recovers the spatial transcriptome of the mouse brain

To demonstrate the potential of STRP-seq to profile anatomically complex tissues, we applied the technique to a coronal section of the mouse brain (Fig. 3). We cryosectioned the brain to obtain five primary and then 679 secondary sections, prepared the cDNA libraries and sequenced them to a depth allowing a transcriptome-wide quantification [8763 genes detected; an average of 40.3 counts per gene per secondary section]. We used this data to verify two key assumptions for the reconstruction. First, we calculated the correlation of the gene expression over the primary sections and confirmed their high similarity (median Pearson’s R = 0.92, Supplementary Fig. 3a). Second, we evaluated the distribution of the noise and found that the read counts closely matched a Poisson distribution at low levels of expression, but that they were overdispersed at higher levels (Supplementary Fig. 3b-c).

**Figure 3.**
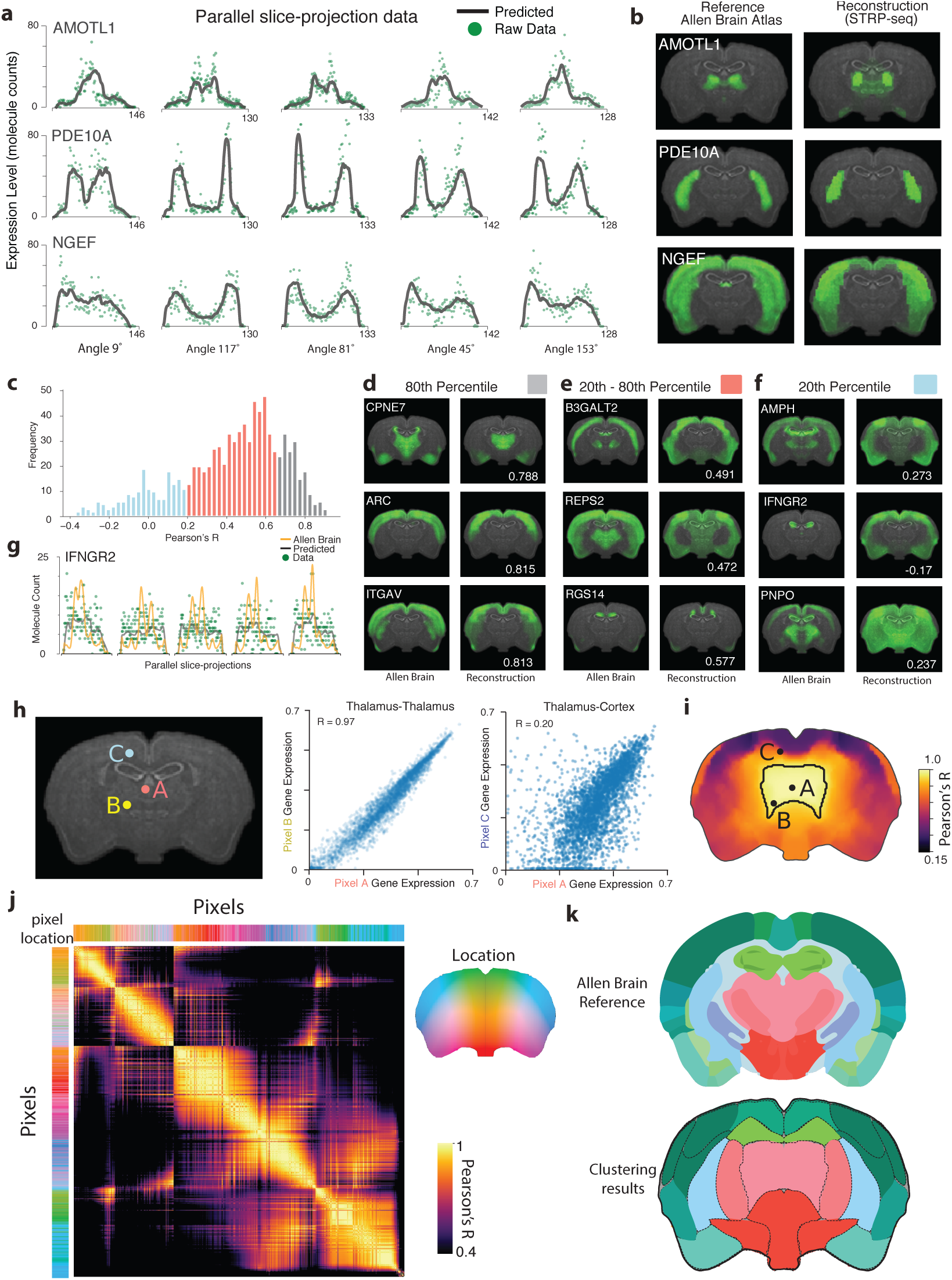
STRP-seq reconstructions of spatial expression profiles for the mouse brain. **a**, Representative examples of parallel-slice projections at different secondary sectioning angles for the genes AMOTL1, PDE10A and NGEF. Raw data points represented by green dots. Parallel-slice projections of the reconstruction as black lines, statistically those values represent the expectation on the counts measured by STRP-seq. **b**, Ground truth ISH data generated by Allen Brain Institute (left) and reconstructed spatial expression profiles (right). **c**, Reconstruction accuracy of STRP-seq as evaluated against the Allen Brain Atlas ISH data. Plot shows the distribution of Pearson’s correlation coefficients. The histogram is color-coded to indicate low accuracy (<20th percentile), intermediate accuracy (20-80th percentile), and high accuracy (>80th percentile). **d**, Examples of high, **e**, intermediate and **f**, low accuracy reconstructions are shown side-to-side with the Allen Brain Atlas ISH data. **g**, Projection of a poorly reconstructed gene (IFNGR2). Ground truth projection depicted in yellow for comparison. **h**, Comparison of the expression profile of the medial thalamus, cortex and ventral thalamus. Markers (A, B, C) indicate different pixels of the image that are used to compute the scatter plots on the right. **i**, Brain section heat map representing the gene expression similarity (Pearson’s correlation coefficient) of each point with respect to point A. Thalamic region outlined by central black contour. **j**, Pairwise correlation distance matrix showing the similarity of gene expression across the pixels of the section. Rows and columns are sorted by similarity using SPIN [37]. **k**, Top: Anatomical annotation by the Allen Brain Institute. Bottom: Unbiased determination of molecular anatomy obtained by pixels clustering. Louvain community detection algorithm was used to cluster the pixels on the basis of their gene expression, no neighborhood constraint was imposed.

Next, we proceeded with the reconstructions (Fig. 3a-b). Based on the results of the previous simulations, we filtered out the genes that were too lowly expressed to allow for reconstructions and retained the parallel-slice projections that varied significantly from the trend of total read counts (3880 genes; see Online Methods Section 4.2). To assess whether we retrieved the correct spatial localization of RNA transcripts, we compared 923 reconstructed genes to the ISH data from the Allen Brain Atlas using Pearson’s correlation coefficient and found a median reconstruction accuracy of R = 0.48 (Fig. 3c; See Online Methods). Notably, gene reconstructions with Pearson’s correlation coefficients as low as 0.2 could already be recognized as similarly distributed to the ground truth, indicating that anatomical localizations can be correctly determined even for low-scoring reconstructions (Fig. 3d-e). In addition to this, across total reconstructions, 2.1% had parallel-slice projections that were incompatible to the Allen Brain Atlas ground truth, suggesting that there might be cases in which ISH and RNA-seq detect different signals, for example, capturing different splicing isoforms (Fig. 3g and Supplementary Fig. 3d).

To further evaluate if STRP-seq profiles are sufficient to delineate anatomical distinctions de novo, we computed pairwise gene expression similarity scores between all the pixels. As predicted, pixels belonging to similar anatomical areas contained more similar gene expression levels. For example, the correlation between two points in the thalamus was 0.97 while it was only 0.20 between points in the thalamus and the cortex (Fig. 3h). Additionally, displaying the similarity between a specific point to the remaining points as an image, we obtain a profile that demarcates different anatomical regions (Fig. 3i,j).

Finally, to actively find those anatomical boundaries suggested by the correlation structure of the data, we clustered pixels using the Louvain community detection algorithm. The clustering output identified regions of neighboring pixels that recapitulate gross anatomical structures defined by the Allen Brain Reference Atlas (Fig. 3k).

Together, these analyses provide evidence that STRP-seq data can be used to reveal the molecular organization of tissue heterogeneity.

### STRP-seq allows for the de novo reconstruction of the molecular anatomy of the lizard brain

To demonstrate the applicability of STRP-seq for examining unconventional or rare samples we profiled brain sections from *Pogona vitticeps*, a non-model organism lizard endemic to semi-arid regions of Australia (Fig. 4a,b; [33], [34]). We used STRP-seq to measure and reconstruct 8,183 annotated genes in the brain of *P. vitticeps* [35]. To validate our approach, we compared a set of images from our reconstructions to ISH experiments and single-cell profiling previously reported, confirming that these genes were enriched in the same locations. (Fig. 4c,d and Supplementary Fig. 4) [34].

**Figure 4.**
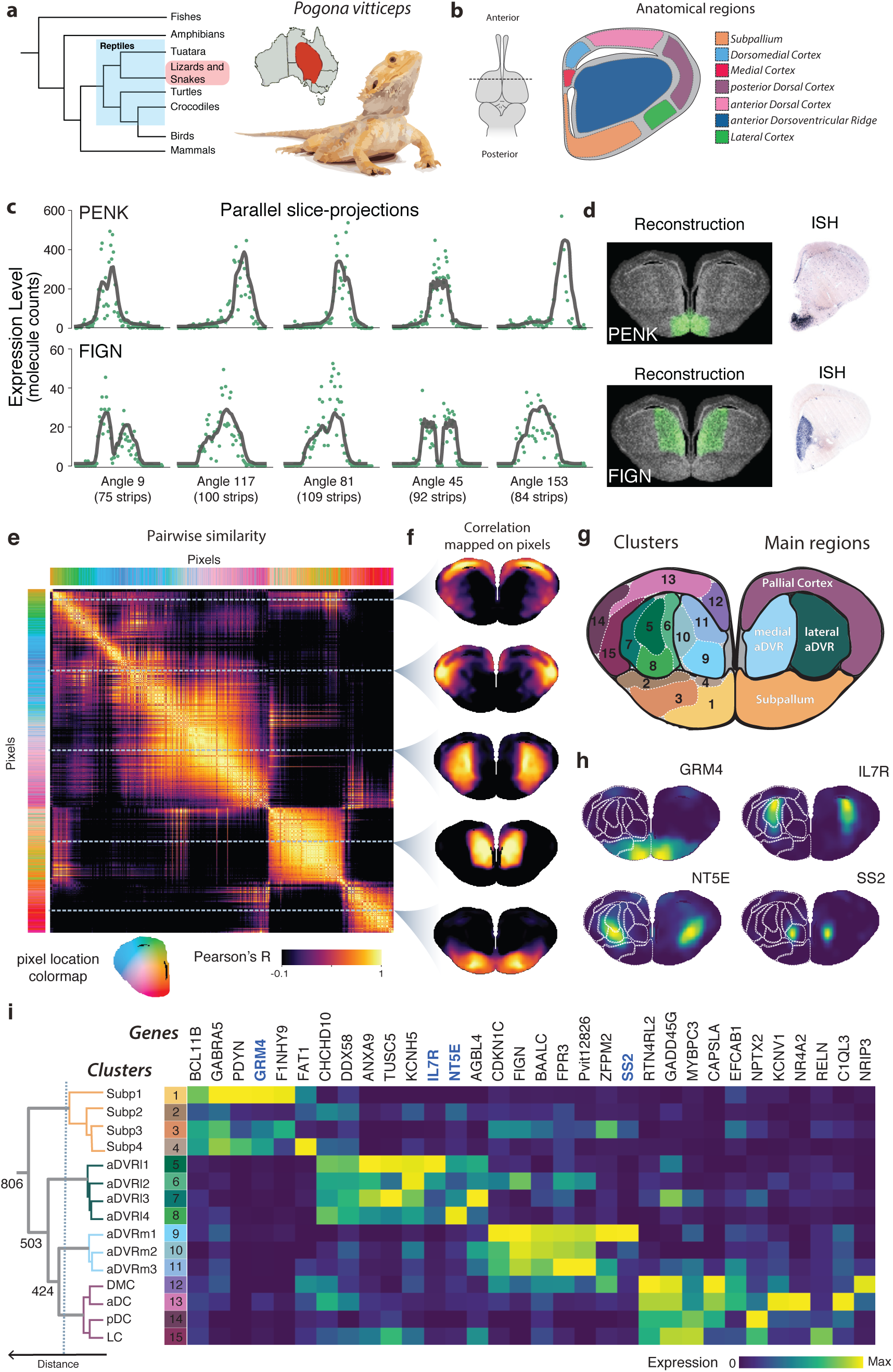
Reconstruction of the molecular anatomy of the lizard brain de novo. **a**, Left: Phylogenetic tree displaying the relation of Lizards to other taxa. Reptilian family highlighted in blue and subsection to which *Pogona vitticeps* belongs in red. Right: Illustration of a *Pogona vitticeps* specimen and its geographical distribution. **b**, Left: Section of tissue in lizard brain that is investigated in our study. Right: A schematic of the anatomical reference annotation for the studied brain section reproduced from Tosches et al. [34]. **c**, Raw STRP-seq data for the genes PENK and FIGN in *P. vitticeps* brain. **d**, STRP-seq reconstruction of expression patterns (left) and validation by ISH (right). **e**, Multivariate analysis of the spatial transcriptome of *P. vitticeps*. The pairwise correlation matrix computed between pixels contains information on the molecular anatomy as highlighted by a block-like structure. The matrix was sorted by the SPIN algorithm. **f**, Selected rows of the matrix are visualized at their anatomical locations the pixel against which the correlation is calculated is marked by the white stripe which runs through the matrix. **g**, Top: the pixels clustered by gene expression reveal the molecular anatomy of the *P. vitticeps* brain. Hierarchical clustering was performed and the tree was pruned at different levels to highlight main clusters and sub-clusters, represented by differently colored regions and by gray lines respectively. Below: Visualization of a selection of genes enriched in distinct sub-clusters. **h**, Gene enrichment score for a variety of genes with respect to individual sub-clusters. **i**, Left: Dendrogram displaying the gene expression distance between the clusters. Numbers indicate the distances between different branches of the tree. Right: Heatmap showing top genes enriched in individual clusters.

According to classical neuroanatomical work, the lizard telencephalon includes the basal ganglia, derived from the ventral telencephalon or subpallium, a layered cortex and the non-layered dorsal-ventricular ridge (DVR), both derived from the dorsal telencephalon or pallium. While the homology of reptilian and mammalian cortices is established, the DVR, which is unique to birds and reptiles, remains an evolutionary dilemma. Recent work, based on single-cell RNA-seq but not spatial transcriptomics, suggested that anterior dorsal-ventricular ridge (aDVR) includes two distinct regions, which have never been recognized in any previous work [36]. To validate this distinction and quantitatively evaluate its relevance in a broader context, we studied the molecular anatomy of the lizard telencephalon using STRP-seq profiles.

First, we performed a multivariate analysis of STRP-seq data examining single pixels. We began by calculating a pairwise gene expression similarity matrix between different locations of the tissue (Fig. 4e). Sorting the similarity matrix using the SPIN algorithm revealed sharply defined groups of covarying pixels [37]. Visualizing the cell-to-cell similarity values in correspondence to the location in the brain, we confirmed that the block-like covariance structure corresponded to anatomical regions in the lizard brain. A more continuous variation was only observed at a finer level within those blocks (Fig. 4f).

To explicitly delineate molecularly distinct regions, we performed unbiased clustering of the pixels (Fig. 4g). We found 15 clusters of contiguous pixels that explained most of the variation present in the data (88.12 %) (Fig. 4g). The 15 clusters provided a nuanced description of the variability (cf. Fig 4e) and could be grouped to identify main regions of the lizard brain. Nonetheless, each cluster was defined by a marked gene expression identity. Gene enrichment analysis highlighted genes that were expressed specifically in individual areas such as SS2, FAT1 and NR4A2 (Fig. 4h,i and Supplementary Fig. 5). Other genes were enriched in multiple clusters, either marking a whole main area (DDX58, BCL11B) or displaying a gradient-like pattern such as IL7R, RTN4RL2 and BAALC (Fig. 4i).

Representing the similarity relations between the 15 clusters using hierarchical clustering revealed a molecular organization of this part of the anterior telencephalon in four main regions. The regions corresponded to the pallial cortex, the subpallium, the medial aDVR and the lateral aDVR (Fig. 4i). The dendrogram suggested a marked dissimilarity between the subpallium and the other regions. Surprisingly, we found that the dissimilarity between the long-believed homogenous medial aDVR and the lateral aDVR was as large as the dissimilarity between the pallial cortex and the lateral aDVR, two functionally distinct regions. When comparing the genes enriched in both regions of the aDVR, we found the lateral part harboring genes encoding for membrane uptake and transport proteins such as ANXA9, TRARG1, KCNH5 and other solute carriers suggesting a possible functional specialization (Fig. 4i and Supplementary Fig. 6a). In contrast, the different regions of the pallial cortex (medial, dorsomedial, anterior dorsal, posterior dorsal and lateral cortex), that have been extensively characterized as functionally distinct, displayed less marked dissimilarities.

### STRP-seq data allows for the analysis of regional identities in the lizard brain

Our clustering analysis of spatial data revealed an unexpected molecular heterogeneity in the lizard anterior telencephalon. To better understand the molecular basis of this difference, we examined the expression pattern of genes characteristic of different neuronal types (Fig. 5a and Supplementary Fig. 6).

**Figure 5.**
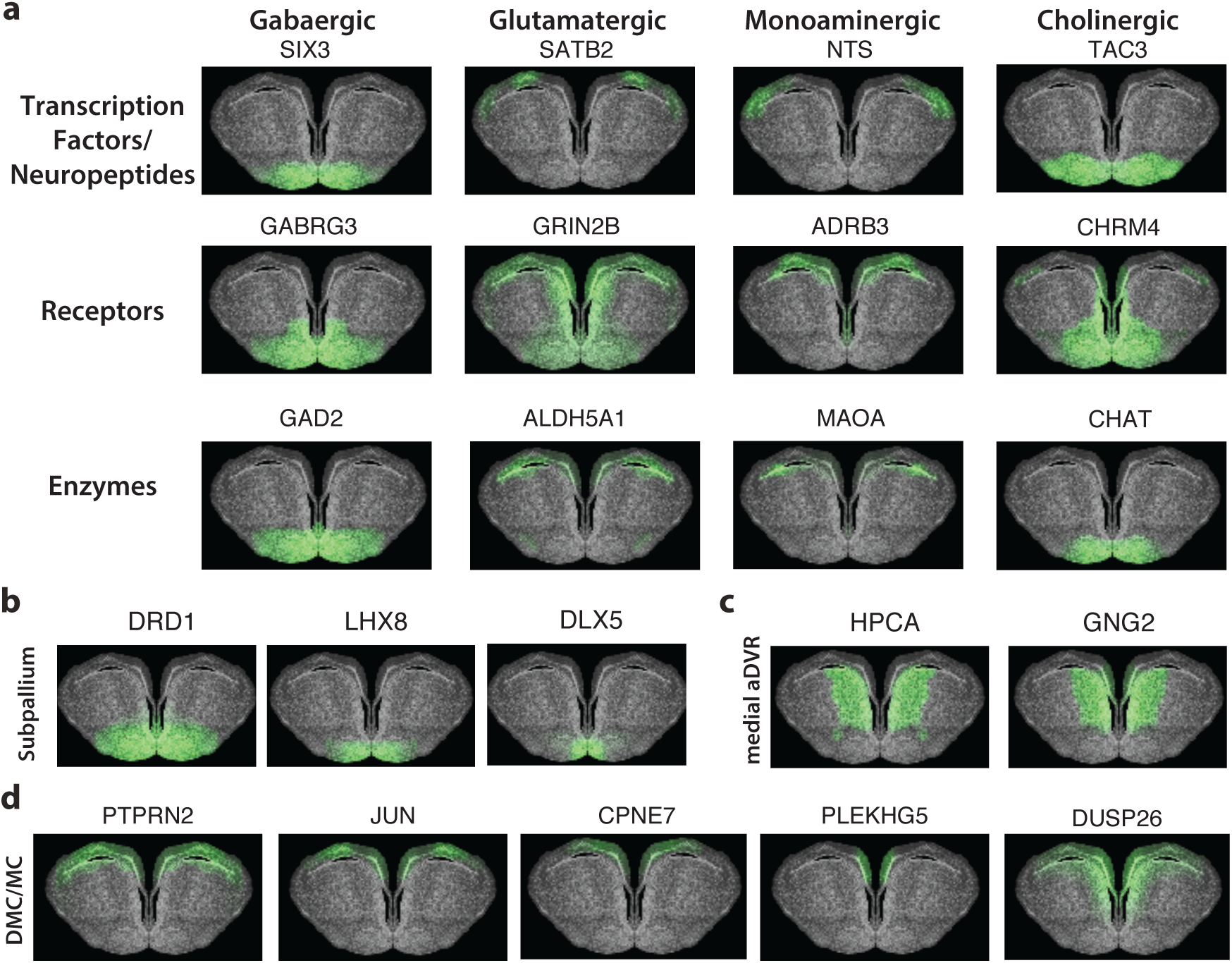
Identification of regional identities in the lizard brain. **a**, Localization of genes belonging to different ontology classes including neuropeptides, transcription factors, neurotransmitter receptors and enzymes involved in neurotransmitter synthesis. Genes are arranged column wise by neurotransmitter type: GABAergic, glutamatergic, monoaminergic (monoamine-receiving neurons) and cholinergic genes. **b**, Visualization of genes expressed in the lizard brain known to be markers of the subpallium (DLX5), the striatal subdivision (DRD1) and the pallidal-preoptic subdivision (LHX8). **c**, Visualization of reptilian medial aDVR marker genes. **d**, Reptilian genes that are homologous of murine hippocampal markers are located in the dorsomedial cortex (DMC) and medial cortex (MC) (PTPRN2: pan-cornus ammonis, JUN: pan-dentate gyrus, CPNE7: granule cells in ventral dentate gyrus, PLEKHG5: granule cells in dorsal dentate gyrus, DUSP26: mossy fibers)

We found genes typically expressed by GABAergic neurons, such as the transcription factors TSHZ1, MEIS2, SIX3 and the postsynaptic GABA receptor GABRG3 expressed ventrally [38]. Additionally, genes expressed by glutamatergic neurons, such as the transcription factors SATB2, NEUROD2 and LHX2 and the postsynaptic glutamate receptor GRIN2B [39], were enriched in ventral and dorsomedial areas. GAD2 and ALDH5A1, which encode for enzymes involved in the biosynthesis of glutamate and GABA, were expressed consistently with the aforementioned markers. In contrast, GOT1, an enzyme involved in the metabolism of glutamate but not in neurotransmission, had a distinct expression pattern (Supplementary Fig. 6b). Furthermore, the genes for the monoamine associated neuropeptide Neurotensin (NTS), the noradrenaline receptor ADRB3, ADRA2A, and the monoamine-degrading enzyme MAOA colocalized dorsally, suggesting that cortical neurons in this area receive monoaminergic inputs [40]. Genes expressed by cholinergic neurons were enriched in ventral regions, in correspondence to the subpallium, including the co-transmitter TAC3, TAC1, the nicotinic acetylcholine receptor CHRM4, and the enzyme CHAT, suggesting the presence of a population cholinergic cells in this region (Fig. 5a and Supplementary Fig. 6b).

To interpret the heterogeneity in a border context, we used the gene expression data to support homology relationships with other vertebrate brain areas. First, we examined in the lizard brain the expression of genes known to characterize the avian and murine subpallium, a region of the vertebrate brain that has an essential role in motoric and sensory functions [41], [42]. We detected marker RNAs of the murine subpallial striatum enriched in the lizard subpallium such as DRD1 and ISL1 (Fig. 5b and Supplementary Fig. 6e). In addition, we localized DLX5 and LHX8, which are known to characterize the entire subpallium and the palladial-preoptic area respectively [41] (Fig. 5b).

Then, we examined the expression of well-described mammalian claustrum markers (RGS12, SYNPR, CNR1, HTR1B, HTR1C, CACNA1B) in the lizard brain and found them to be expressed in the medial aDVR, where they colocalized with bona fide markers for this region (HPCA, GNG2, ADARB2, RORB, CPNE4) (Fig. 5c and Supplementary Fig. 6c,d). This finding confirms that the medial aDVR in lizards is the claustrum homologue, as suggested by electrophysiological characterization, axonal tracing and single-cell RNAseq [36],[43].

Lastly, to confirm the hypothesis that the reptilian pallial cortex contains a region homologous to the hippocampus, we localized genes that are known to be expressed in different subregions of the murine hippocampus [34], [44]. Expressed in the medial and dorsomedial cortex we found markers of the cornus ammonis (CA) region 1-4, genes characterizing hippocampal mossy fibers and markers of granule cells in the ventral and dorsal dentate gyrus (Fig. 5d and Supplementary Fig. 6f).

Finally, we performed an independent comparison of the examined region with mammalian data, by both cross-referencing the STRP-seq data with a recently proposed molecular taxonomy of cell types in the mouse brain and scoring the correspondence between the expression pattern of a given pixel and of the cell clades (See Online Methods; Supplementary Fig. 7; [45]). Importantly, the analysis confirmed the localization of the dentate gyrus in the medial cortex and the ventral positioning of the subpallium (Supplementary Fig. 7). Interestingly, for the anterior dorsal-ventricular ridge, we found a set of genes compatible with telencephalic inhibitory neurons and a non-cholinergic motor neuron signature in the medial and lateral aDVR respectively. Moreover, we observed an enrichment of astrocytic markers in the ventricular area, most likely marking reptilian ependymoglial cells that surround the lateral ventricle (Supplementary Fig. 7; [34]).

## DISCUSSION

Here, we present a versatile framework that allows to transform existing low-input NGS techniques into methods capable of encoding spatial information through compressed sampling and a probabilistic image reconstruction algorithm [27]. We report an implementation of our framework for studying transcriptomic data, STRP-seq, and demonstrate its application to profiling rare samples such as the brain of the Australian lizard *P. vitticeps*. To facilitate the adaptation of the presented framework to applications beyond gene expression studies, we provide *Tomographer*, a versatile software package to solve fast and parallelized image reconstruction problems for genomic data.

One limit of our approach is its relatively low resolution when compared to imaging based methods and its inability to reveal discontinuous, checkboard-like patterns [10]. For example, our method does not allow for discrimination of the signal at the single-cell level. Instead, it is designed to capture broader tissue-level patterns of expression. For these reasons, we envision the method to be most useful as a way to link single-cell resolved data to its spatial context, to profile unique samples (e.g. biopsies), and to study the molecular anatomy of non-model organisms.

We demonstrate how STRP-seq can strengthen our understanding of evolutionary relationships between brain regions. We show this by supporting hypotheses on region homology with molecular data and by presenting further evidence for an anatomically divided aDVR [34], [36]. In particular, we suggest a starker molecular distinction than previously thought. In the medial aDVR, which is considered the reptilian claustrum homologue, we discovered a gene expression signature related to mammalian telencephalic inhibitory neurons, suggesting a differential recruitment of this set of genes to glutamatergic cells of the medial aDVR. In the lateral aDVR, we unexpectedly detected the expression of genes related to non-cholinergic motor neurons, suggesting that we capture mRNA of neurons projection from the brain stem to the lateral aDVR [46].

Taken together, we provide a fundamental generic profiling scheme compatible with different types of genomics measurements that can be set up at a low cost and without the need of specialized instrumentation. We envision our framework will help bridge the gap between the various available NGS techniques and their spatial counterpart.

## ACKNOWLEDGEMENTS

We thank Sten Linnarsson (Karolinska Insitutet) for allowing proof of principle tests in his laboratory, Noam Shental, Bar Shalem (Open University Israel) and Amit Zeisel (Technion) for stimulating discussions, Markus Schuelke (Charité) for enabling the participation of C.G.S. in this project. This work was supported by a grant from the Swiss National Science Foundation (CRSK-3_190495) to G.L.M.

## AUTHOR CONTRIBUTIONS

G.L.M. conceived the study design and supervised the project. G.L.M. H.H.S. and C.G.S. analyzed, annotated and interpreted the tomography data and wrote the manuscript. M.A.T. and T.J. performed *P. vitticeps* experiments, in situ hybridization and contributed interpreting the results together with G.L.. G.L.M., A.R., H.S designed and wrote the reconstruction algorithm. S.C. and L.E.B. designed the cryosectioning scheme. S.C. and J.S. performed the mouse experiments and sectioning. P.L. ran the bioinformatics pipeline. F.P.A.D. built the companion website. All the authors critically reviewed the manuscript and approved the final version.

## CODE AND DATA AVAILABILITY

Source code and templates for customized 3D-printable cryosectioning adaptor manifolds are available at https://github.com/lamanno-epfl/tomographer. RNA-seq data is available at the Gene Expression Omnibus repository (GEO; https://www.ncbi.nlm.nih.gov/geo) under accession GSE152989. Results of the reconstruction can be accessed at this paper companion website https://strpseq-viewer.epfl.ch/.

---

**Supplementary Figure 1.**
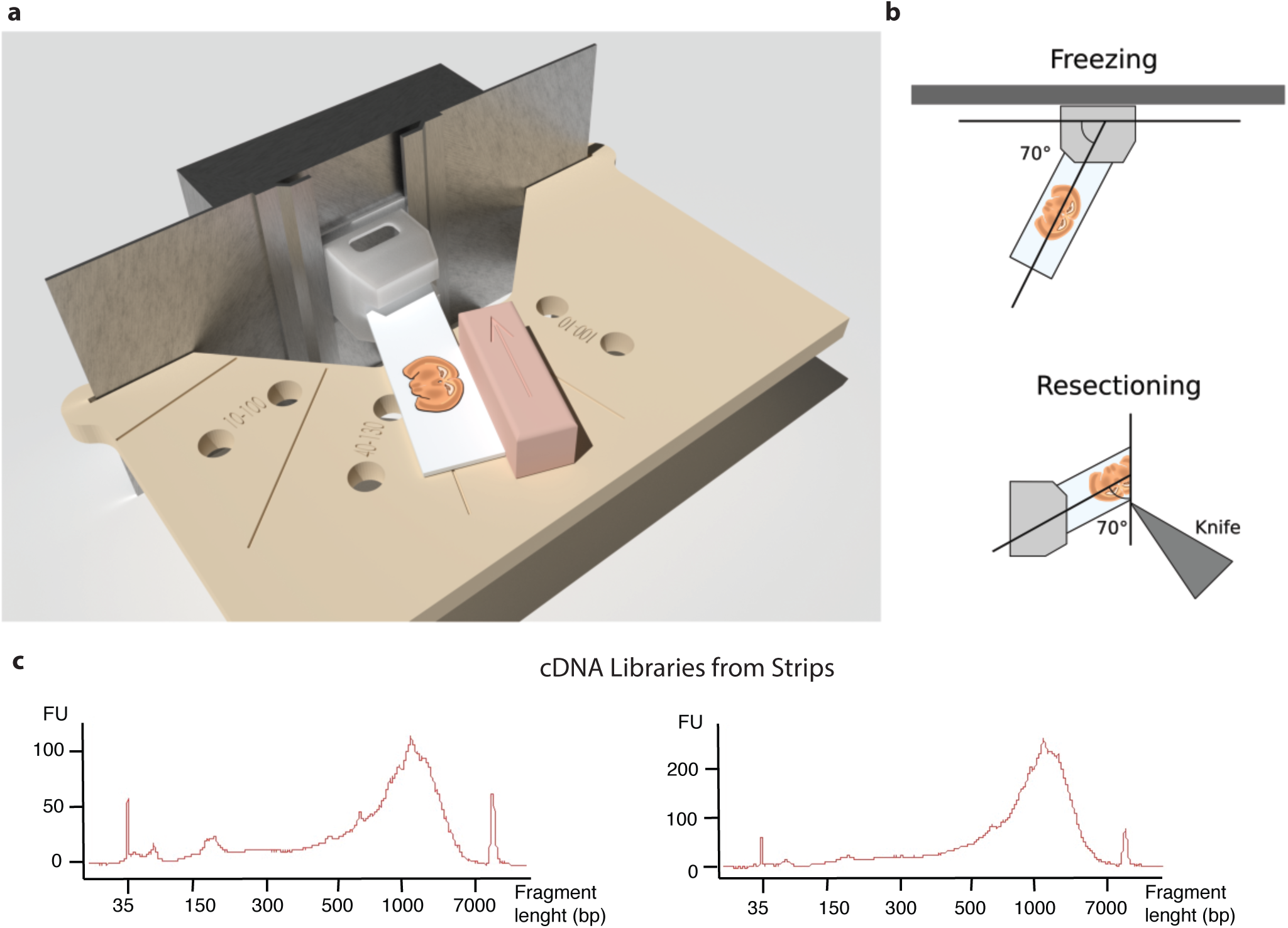
Generation of secondary sections and cDNA libraries. **a**, Representation of the device used for positioning primary sections to obtain secondary sections. The primary section is mounted on a slab of ice and positioned in a block according to a predefined angle using a 3D printed manifold. **b**, The slab is frozen at the predefined angle with the base (top). The base is mounted in a cryostat and sliced at a constant width to generate strips at the desired angle (bottom). **c**, Example of representative bioanalyzer profiles of the cDNA library produced from two strips.

**Supplementary Figure 2.**
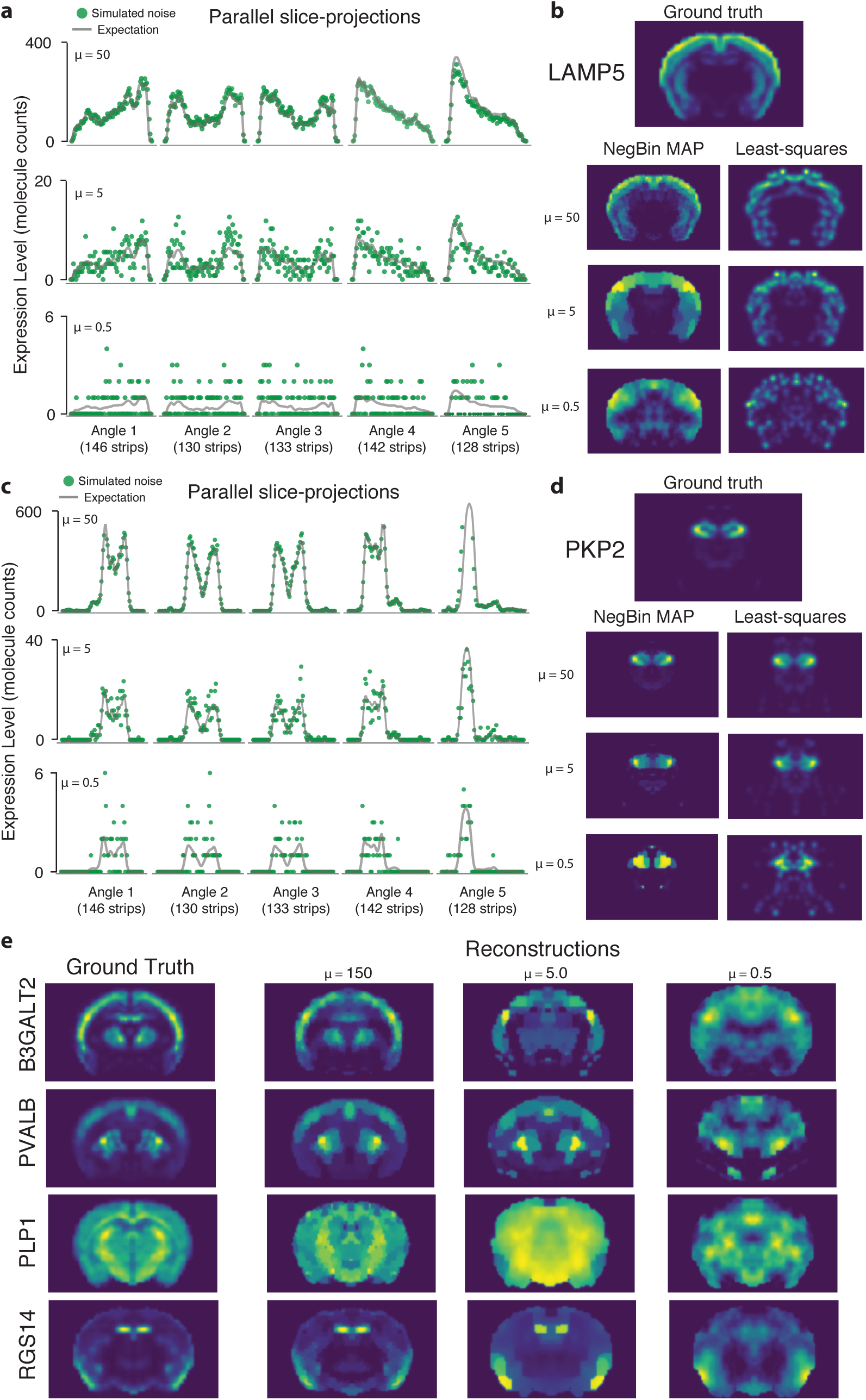
*Tomographer*’s reconstructions at different signal-to-noise ratio conditions. **a**, Parallel-slice projections of simulated data overlaid with expectation for LAMP5 for three degrees of noise. **b**, LAMP5 mathematical image reconstructions using Negative Binomial maximum a posteriori estimation and least-squares approach constrained to positive results. **c**, Parallel-slice projections of simulated data overlaid with expectation for PKP2. **d**, PKP2 mathematical image reconstructions using Negative Binomial Maximum A Posteriori Estimation and least-squares approach constrained to positive results. **e**, Ground truth and reconstructions using Negative Binomial maximum a posteriori estimation for B3GALT2, PVALB, PLP1 and RGS14.

**Supplementary Figure 3.**
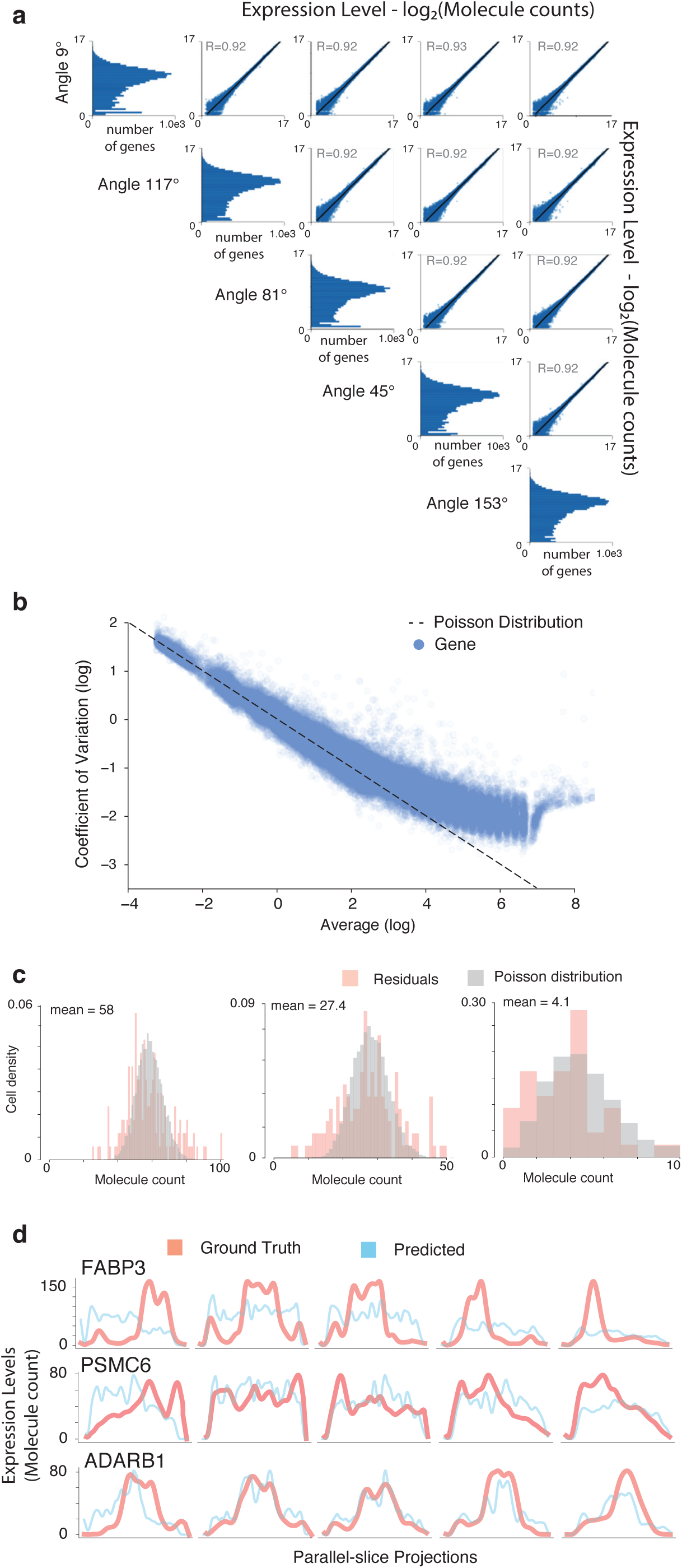
Verification of the assumptions used in the *Tomographer*’s algorithm. **a**, Scatterplot matrix comparing the total gene expression of the primary sections (sum of the counts sequenced across all the secondary sections for each angle). Pearson’s R is indicated for each comparison. Histograms on the diagonal show the total molecule count distribution across genes. **b**, Evaluation of the over-dispersion of the count distribution for different genes across the parallel-slice projections. Scatterplots show the coefficient of variation as a function of the mean; the relationship expected for a Poisson distribution is displayed as a reference. Note that in the scatter each gene appears multiple times: since genes are detected at different levels across the parallel-slice projection, we group points by the reconstruction expectation and, for each group, we compute the coefficient of variation and the mean and plot a dot (See Online Methods). **c**, Molecule count distribution for high, medium and low gene expression. In red, the histogram of observed counts compared with, in gray, an expected Poisson distribution with the same mean. **d**, Selection of genes with low, medium and high correlation values between provided data and Allen Brain ISH projection. Ground truth projection (red) and predicted projection (blue) visualized.

**Supplementary Figure 4.**
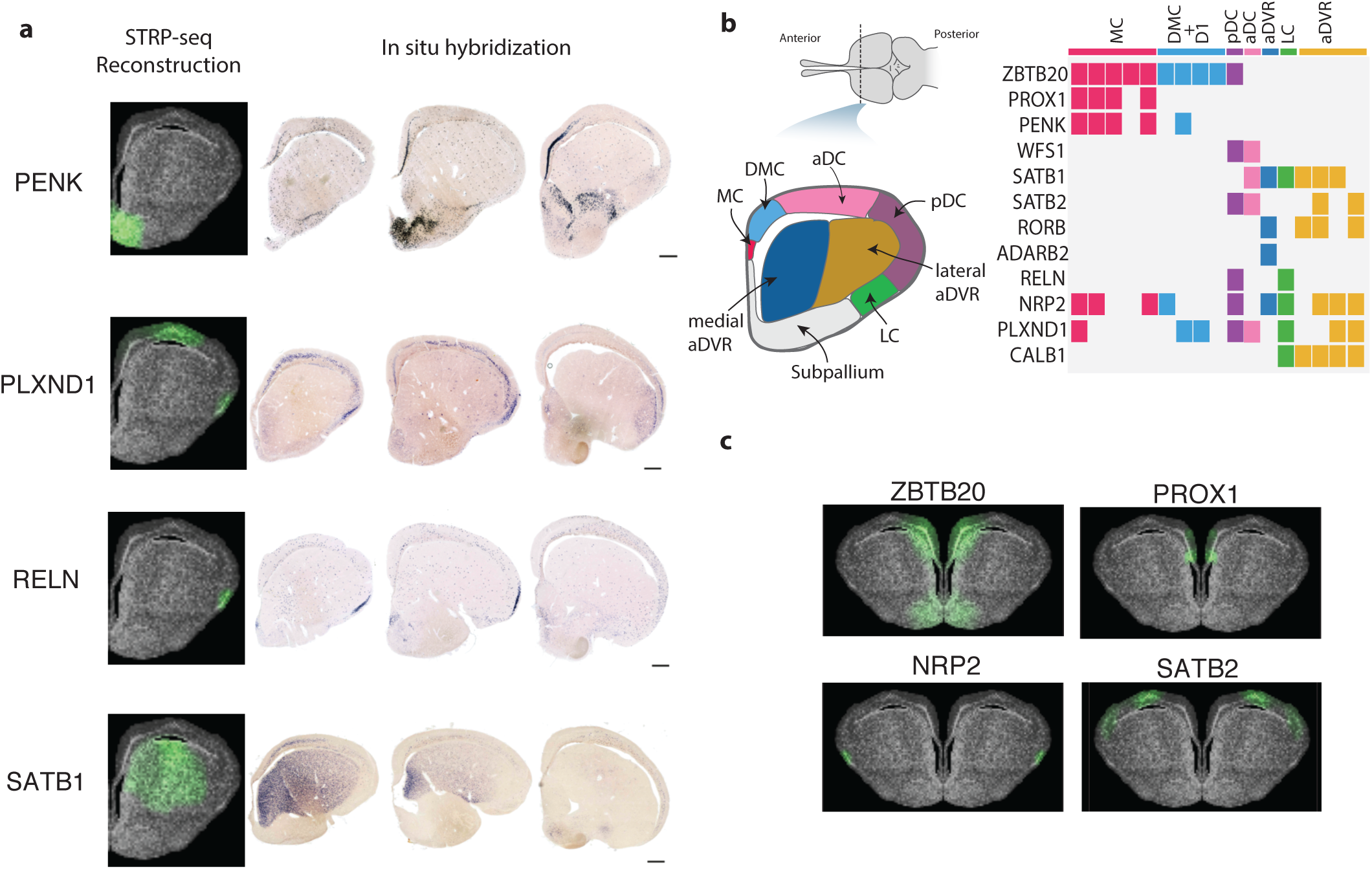
Validation of STRP-seq reconstructions in lizard. **a**, Left: STRP-seq reconstruction. Right: ISH staining for selected genes across different sections in the reptilian brain taken from Tosches et al. [34] (MC: medial cortex, DMC: dorsomedial cortex, D1: dorsal cortex area D1, aDC: anterior dorsal cortex, LC: lateral cortex, aDVR: anterior dorsal-ventricular ridge) **b**, Proposed partitioning of the reptilian brain tissue according to [34] (left) and a binarized summary of the gene expression profile of cell types identified by single-cell RNA-seq grouped by their putative localization of various subregions (right). **c**, *Tomographer* reconstructions of the genes in *P. vitticeps* corresponding to the genes displayed in (b) [34]. The genes not displayed were already shown as part of another other display figure element (cf. Fig. 4, 5, Suppl. Fig. 5).

**Supplementary Figure 5.**
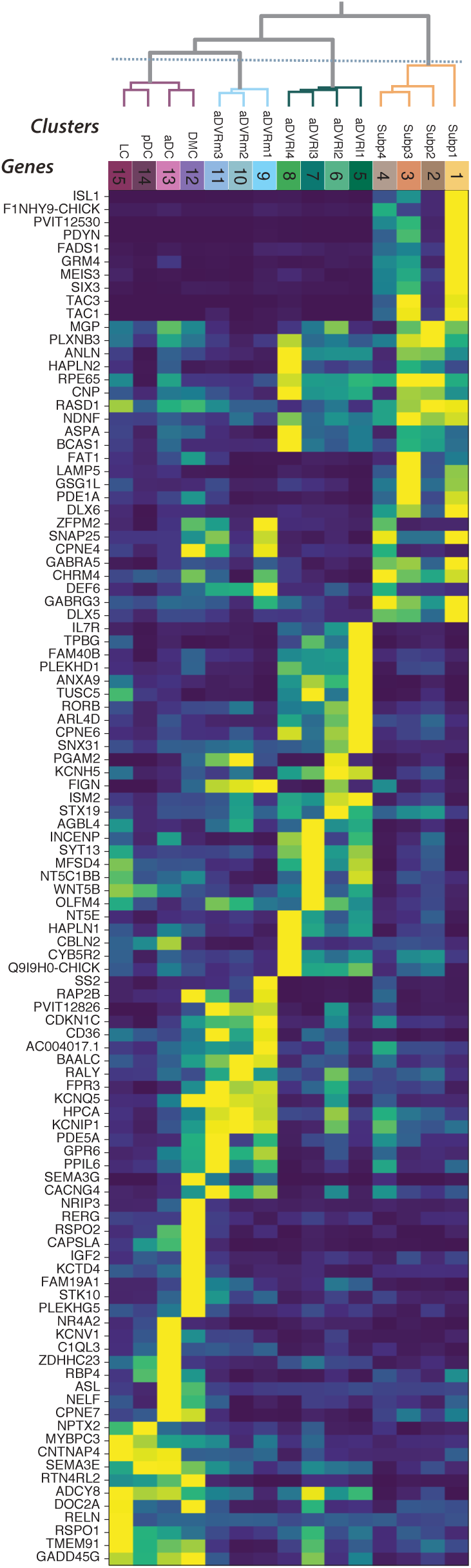
Genes enriched in clusters of the lizard brain. Heatmap displaying genes enriched in each of the clustered areas. Genes are sorted by the cluster they are maximally expressed in.

**Supplementary Figure 6.**
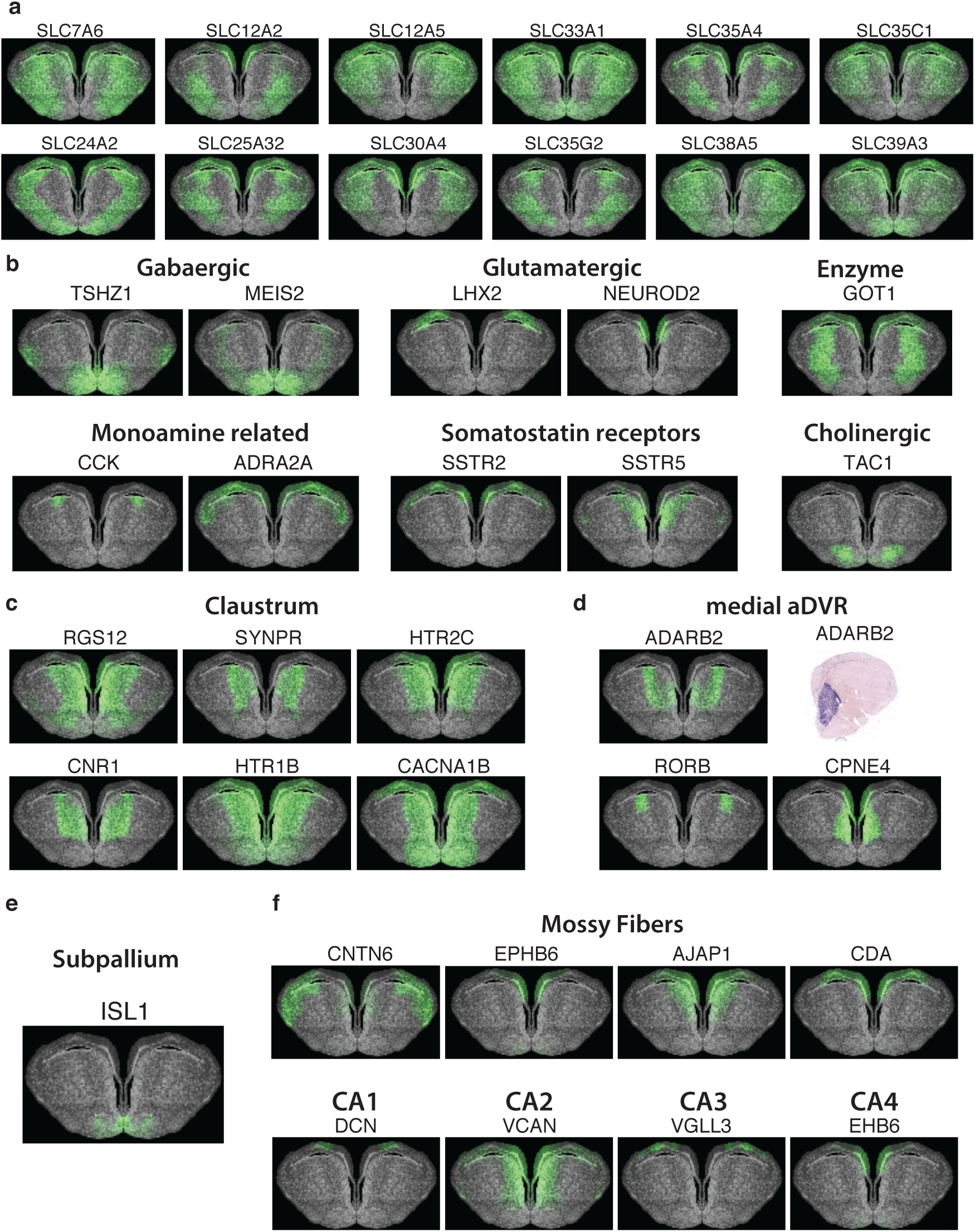
Genes in the lizard brain by cell type and regional specificity. **a**, Genes encoding for transporter proteins from the solute carrier (SLC) group enriched in the lateral aDVR. **b**, STRP-seq reconstructions of cell-type specific genes. GABAergic transcription factors TSHZ1, MEIS2 enriched in ventral regions, glutamatergic transcription factors LHX2 and NEUROD2 enriched dorsally. The glutamate metabolizing, neurotransmitter-non-specific enzyme GOT1 showing a distinct expression pattern from enzymes involved in transmitter synthesis of glutamatergic and GABAergic neurons (cf. Fig. 5a). The monoamine associated co-transmitter CCK, noradrenaline receptor ADRA2A and somatostatin receptors SSTR2, SSTR5 are enriched in dorsal regions. Cholinergic co-transmitter TAC1 enriched in the subpallium. **c-d**, Markers of the murine claustrum colocalize with lizard medial aDVR marker genes. Inset in (c) shows an ISH validation for ADARB2. **e**, A marker gene of the murine subpallial striatum (ISL1) expressed in the lizard subpallium. **f**, Marker genes of murine mossy fibers and the CA regions enriched in the dorsal and dorso-medial cortex of the lizard pallium.

**Supplementary Figure 7.**
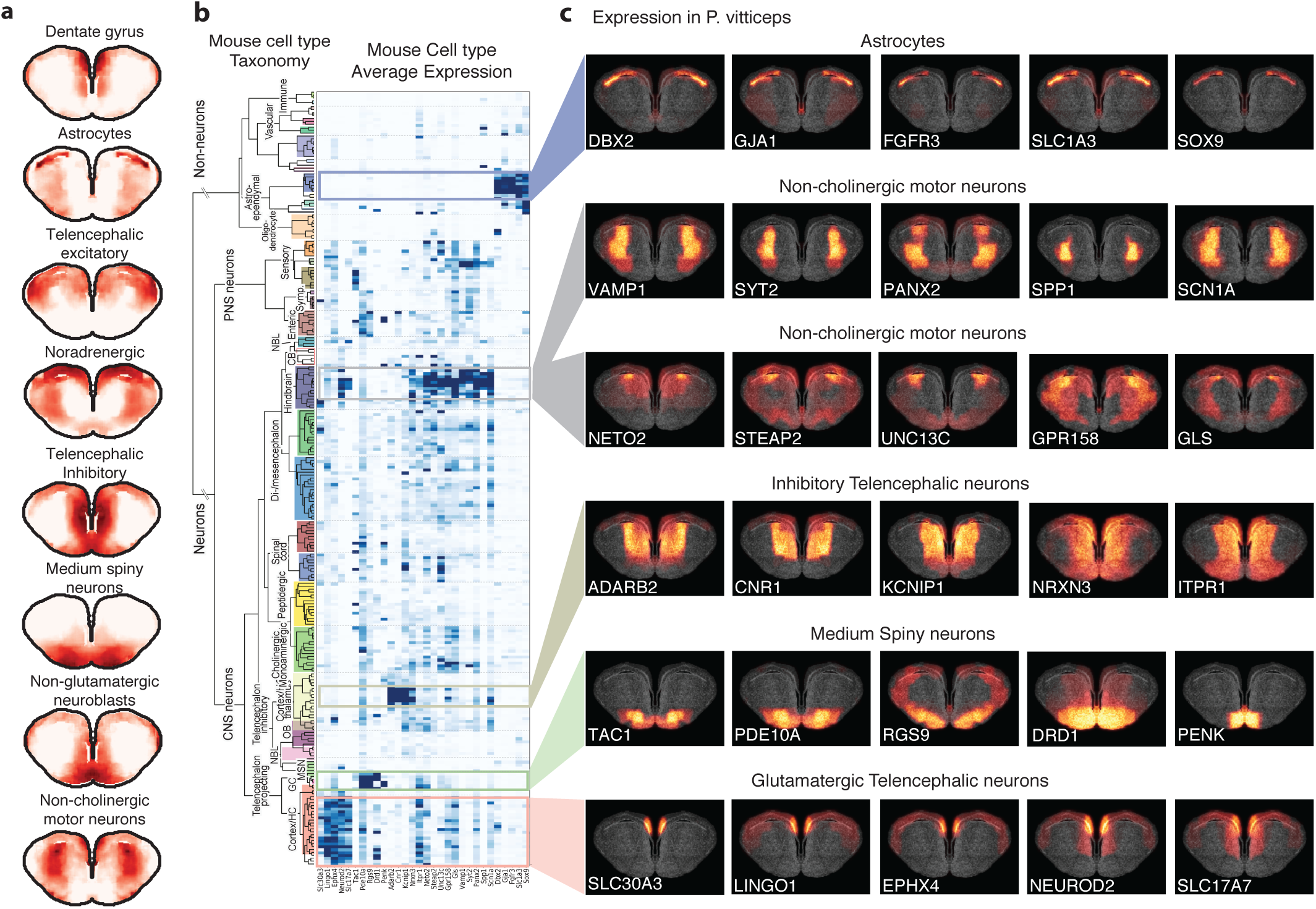
Localization of cell-type signatures in the brain of *P. vitticeps*. **a**, Results of a dictionary learning analysis where the full gene expression pattern of each brain location is approximated by a linear combination of mouse cell type expression patterns from [45] (See Online Methods). The figure shows the coefficients computed for each cell type category: the darker the color the more important was the category to explain the observed gene expression pattern. Categories are displayed in descending order according to the coefficient peak. Medium Spiny Neurons marking the subpallium (striatum), Glutamatergic Telencephalic neurons marking the medial cortex (dentate gyrus) **b**, The Zeisel Mouse cell type taxonomy [45] flanked by a heatmap displaying the average gene expression in correspondence to each gene. Genes displayed are a selection of marker genes that colocalized in a lizard region and one or more cell-type clades. **c**, Visualization of the gene expression pattern found in *P. vitticeps* tissue. The genes are the homologues ones displayed by the heatmap on the left.

## Online Methods

### 1 Experimental Work

#### 1.1 Animal work

The CD-1 mouse was obtained by Charles River (Germany). Mice were housed in rooms with a regular dark/light cycle and fed a standard rodent diet and water ad libitum. All animal procedures were approved by the local ethics committee in Stockholm (Stockholms djurförsöksetiska nämnd) and followed the Directive 2010/63/EU of the European Parliament and of the Council, the Swedish Animal Welfare Act (Djurskyddslagen: SFS 1988:534), the Swedish Animal Welfare Ordinance (Djurskyddsförordningen: SFS 1988:539) and the provisions regarding the use of animals for scientific purposes: DFS 2004:15 and SJVFS 2012:26. *P. vitticeps* experiments were conducted at the Max Plank Institute for brain research and followed the Hessian, German and EU laws on animal experimentation. Lizards were sacrificed according to § 4 (3) Tierschutzgesetz (TierSchG, German animal welfare law) and § 2 Tierschutz-Versuchstierverordnung (TierSchVersV).

#### 1.2 Cryosectioning

Agarose membranes for cryosectioning were prepared by allocating a drop of 3% low melting temperature agarose stained with Orange-G (Sigma) between two superfrost glass slides (Thermo). To remove excess agarose, a metal frame from a laser capture microdissection slide (LCM0521, Thermo) was placed on the membrane and used as a trimming guide. Fresh frozen and OCT (Tissue-Tek) embedded brains were cut into 14 µm thick sections and collected on the agarose membrane which was trimmed once again. A metal frame was placed on the prepared slide and the created well was filled with OCT and frozen on dry ice. The glass slide was then replaced with a second metal frame which was filled with OCT and frozen again. To cut the collected sections at a predefined angle, each sample was removed from the metal frame using a scalpel and mounted onto a customized positioning device (Suppl. Fig 1a). The device consists of a metal wall that can be attached to a cryomold and a sample retainer that can position the tissue slaps at a predefined angle relative to the metal wall using a mobile guide. Before each use the positioning device was placed on dry ice and the right cutting angle was selected by moving the mobile guide into a respective hole of the sample table. The cryomold was attached to the positioning device by cutting one of its plastic walls with a scalpel and sliding it into the clip of the metal wall so that the rectangular hole aligns with the surface of the table. It was then filled with Optimal Cutting temperature compound (OCT) up to the level of the slit and frozen rapidly using a head sink mounted behind the metal wall. To start the sectioning process, a tissue slab was placed on the sample retainer, aligned with the angle guide and slit into the cryomold. The sample was frozen in the OCT block at the correct angle and repositioned on the cryostat. 70 µm strips were sliced, collected with forceps and transferred in a 96-wells plate kept inside the cryostat. The angle of the cryostat sample holder was not changed between samples. All samples were collected the same day and stored at -80°C.

#### 1.3 Preparation of cDNA library

We prepared poly-T coated magnetic microbeads incubating for 15min 24 µL (per plate, 0.25 µL per sample) of MyOne C1 streptavidin beads washed using a magnet and resuspended in Binding Washing Tween (BWT) 1x buffer (5 mM Tris-HCl, 1 M NaCl, 0.5 mM EDTA, 0.1% Tween 20) containing 5M STRTC1 P1A T31 Primer (5’Biotin-AAT GAT ACG GCG ACC ACC GAT CGT TTT TTT TTT TTT TTT TTT TTT TTT TTT TTT). The beads were then washed twice in Lithium Washing Tween (LiWT) buffer (10 mM Tris-HCl pH 7.5, 0.15 M LiCl, 1 mM EDTA) resuspended in 480 µL (per plate, 5 µL per sample) of the same buffer, ready to be added to the lysate. Plates containing the cryosectioned strips were quickly spun down and placed on a thermally conductive metal block positioned on dry ice. To lyse the strips 37 µL of Lysis/Binding buffer (LBB) (100 mM Tris-HCl, pH 7.5, 500 mM LiCl, 10 mM EDTA, 1% Lithium dodecyl sulfate (LiDS), 5 mM dithiothreitol (DTT)) were added to each well of the plate, 0.4 µL of 1:1000 ERCC per sample (corresponding to 24.8M molecules) were also added as an internal control. The plate was maintained on dry ice and the buffer was allowed to freeze. The plate was then alternatively vortexed for 40 s and briefly centrifuged for a total of 3 times. The plate was then incubated at 37°C for 7 minutes. Vortex agitation and spinning down centrifugation were repeated twice and then 5 µL of the previously prepared mix containing the poly-T coated beads were added. The samples were incubated 2 min at 70°C on a thermal cycle, were let equilibrate at room temperature and incubated 30 min at room temperature. The beads were bound to a magnet and the samples washed with LiWT twice each time carefully resuspending the beads by vortexing. Finally beads of each well were resuspended in 7.5 µL of RT Mix and carefully resuspended by pipetting. RT Mix was prepared mixing the following reagents: 163 µL of 5X SuperScript First-strand buffer (18064-014 Thermo), 4.9 µL of 1M MgCl2, 135 µL of 5M Betaine, 48 µL dNTPs, 20 µL DTT 0.1M, 7 µL 10% Tween 20, 20 µL RNAse inhibitor (Takara), 50 µL Template Switching Oligo 40 µM (C1-P1-RNA-TSO: 5’Biotin-rArArU rGrArU rArCrG rGrCrG rArCrC rArCrC rGrArU rNrNrN rNrNrG rGrG), 40 µL Superscript II (18064-014 Thermo), 349 µL of RNAse-free water. The plate was incubated for 1 h 30 min at 42°C and 10 min at 72°C the beads were resuspended by vigorous agitation of the plate every 15 minutes. The beads were bound using a magnet and supernatant discarded. The beads, coated with cDNA, were then resuspended in 16 µL PCR mix and the following PCR program was run for a total of 18 cycles. 3 min at 95°C; 5 cycles of: 20s at 98°C - 4 min at 62°C - 6 min at 72°C; 9 cycles of: 20s at 98°C - 30s at 68°C - 6 min at 72°C; 4 cycles of: 20s at 98°C - 30s at 68°C - 6 min at 72°C; 10 min at 72°C; Hold at 4°C. The PCR mix was assembled by combining the following reagents: 1 mL of 2x KAPA Hot Start Mix, 80 µL of 5M Betaine, 2 µL 10% Tween 20, 80 µL of PCR primer 10 µL (C1-P1-PCR-2: 5’Biotin-GAA TGA TAC GGC GAC CAC CGA T) and 838 µL RNAse-free water.

#### 1.4 Library preparation and sequencing

Samples were diluted to 2 ng/µL of PCR product and a random sample of wells was inspected on an Agilent Bioanalyzer using a high sensitivity DNA kit. The first and last secondary section for which a cDNA library was clearly visible on the bio-analyzer were identified and a buffer of 6 extra samples before and after were retained for sequencing. Sequencing library was prepared by “tagmentation”, that is by simultaneously fragmenting and barcoding using Tn5 DNA transposase [1]. Briefly, 6 µL of amplified cDNA were transferred to a multi-well plate containing barcoded adaptors. Tagmetation buffer (6 µM TAPS-NaOH, pH 8.5, 25 mM MgCl2 and 50% DMF) and transposome stock were added before incubating the plate at 55 C for 5 min. The fragmented cDNA products were bound with an excess of Dynabeads MyOne Streptavidin C1 beads in order to retain only the 5- and 3-most fragments. All fractions were pooled, the beads were immobilized, washed and 3 fragments cleaved with a PvuI restriction reaction. Finally, the single-stranded library was eluted in water. Samples were sequenced on a Illumina HiSeq 2000 at a total depth of 1 billion reads per experiment for both the mouse and lizard experiment.

#### 1.5 Read mapping pipeline

To obtain the table of UMI counts corresponding to each strip we processed the sequencing with a custom pipeline that is thoroughly described elsewhere [2]. The pipeline was designed and optimized for the STRT-seq chemistry from which we adapted our RNA-seq protocol. Briefly, the sequencing data was aligned to the mouse genome using Bowtie, different quality controls were applied. The parameter *filter-singletons* was set to False since we expected more real molecules to be supported by single UMI read than in a single-cell scenario. For *Pogona vitticeps* we used the same pipeline but, in absence of a high quality genome, we performed the mapping directly on an assembled transcriptome reported by Georges et al., 2015 [3]. The assembled transcript models were extended at the 5’ by adding the 200bp upstream determined using genome contigs to buffer that inaccurate TSS definition.

### 2 Models and Computational Approaches

#### 2.1 Nomenclature

- The two-dimensional expression pattern for each gene can be represented as a matrix **X** ∈ ℝ^*n,m*^. We denote the intensity of pixel at position *i, j* with *X*_*i,j*_. We also introduce an alternative notation for the same object. **x** ∈ ℝ^*nm*^ is the vector that is obtained by flattening **X**, that is: *X*_*i,j*_ = *x*_*m*(*i-*1)+*j*_. These two notations are used alternatively in different parts of the text to make formulations easier to interpret.
- For each gene, we indicate data from each parallel-slice projection over a given angle γ with a vector 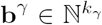 where *k*_*γ*_is the number of strips obtained cutting at angle γ so that each entry 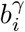 contains the read count measured in correspondence to the *i*^*th*^ strip. Henceforth we will omit the γ indices and refer to the entire set of parallel-slice projection with the stacked column vector **b** = (**b**^*αT*^, **b**^*βT*^, *…*, **b**^*ωT*^)^*T*^. Instead, we use 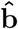 to indicate the parallel-slice projection vector predicted by the model.
- The information regarding the orientation and width of the slices is contained in a matrix **A** ∈ ℝ^*k,nm*^ which we refer to as the design matrix. **A** is a sparse matrix that is constructed so that *A*_*ij*_ is set to 1 if the entire pixel *x*_*j*_ was sampled by the *k*^*th*^ strip **b**_*k*_, to a fraction if only part of the pixel was sampled, zero otherwise (details in Section 2.4). The design matrix can be thought as a linear operator that acts onto the image (i.e. the vectorized **x**) to return the expected molecule count the secondary slice-projections that is 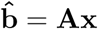.

#### 2.2 Assumptions

- The pixel intensities and differences of pixel intensities are assumed to be drawn from Exponential and Laplacian distributions, respectively.
  – Genes are assumed to be relatively localized and have pixel intensity histograms that accumulate on low values. We place an exponential prior on pixel values *x ∼* Exp(,*λ* = α).
  – The signal we want to reconstruct appears as an image. This implies that transitions from pixel to pixel are generally smooth and only rarely sharp (at edges of gene expression). We therefore impose a Laplacian prior on both dimensions of the pixel gradient. In other words, we consider the two priors (*x*_*i*+1,*j*_ *x*_*i,j*_) *∼* Laplacian(*µ* = 0, 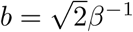) and (*x*_*i,j*+1_ *x*_*i,j*_) *∼* Laplacian(*µ* = 0, 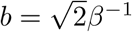).
- Each gene’s expression profile is independent from another.
- Each angle of the parallel-slice projection *b*_*γ*_ is a re-sampling of the same distribution in ℝ^2^.
- The sampling distribution of the parallel-slice projection **b** given the pixel intensity values follows a Negative Binomial Distribution whose mean is given by 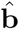.

#### 2.3 Formulation of the optimization problem

The objective is to estimate the pixel intensity values given the values of the observed molecule counts, **b**. The task is framed as a maximum a posteriori estimation problem where we posit that the sampling distribution of the parallel-slice projection **b** given the pixel intensity values follows a Negative Binomial Distribution whose mean and variance are 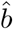 and 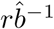, respectively.

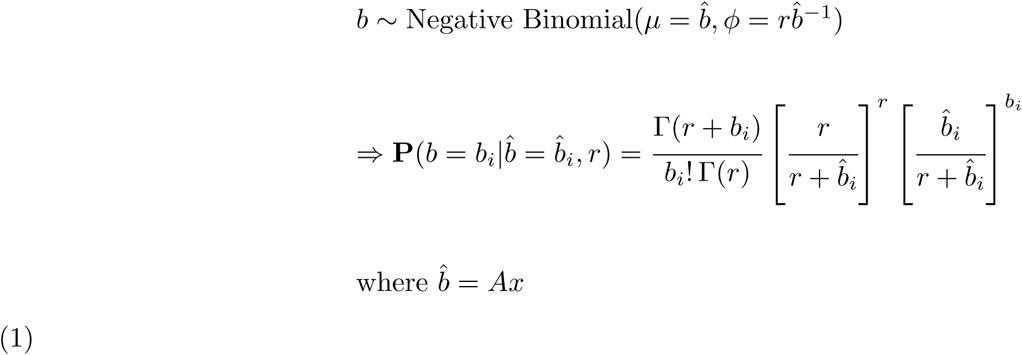

The pixel intensities and difference of pixel intensities are assumed to be drawn from an Exponential and Laplacian distribution, respectively.

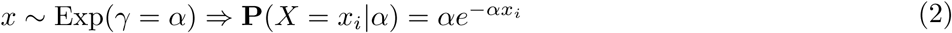

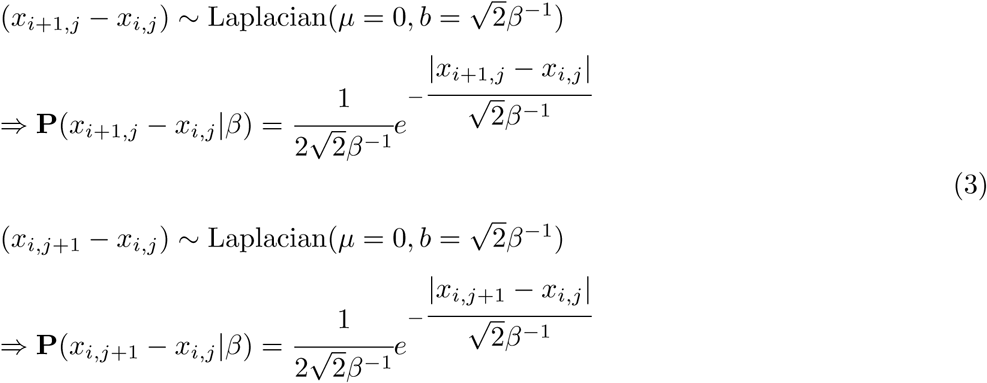

Deriving the posterior distribution of the pixel intensity values **x** using Bayes Theorem gives us

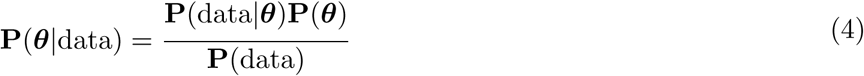

Since **P**(data) is our vector containing the experimentally obtained values, it is a constant and therefore independent of the parameters to be estimated. Thus, it can be ignored to simplify the expression to

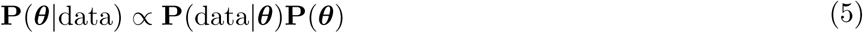

Incorporating our model and its distributions as defined above we can then write

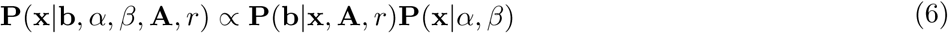

The maximum a posteriori estimate for the pixel intensity values *x* becomes

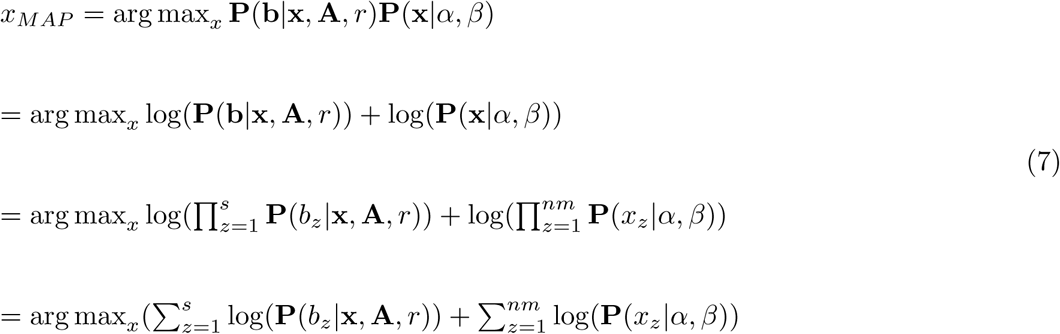

Substituting the probability distributions defined previously, the *x*_*MAP*_ expression can be written as follows:

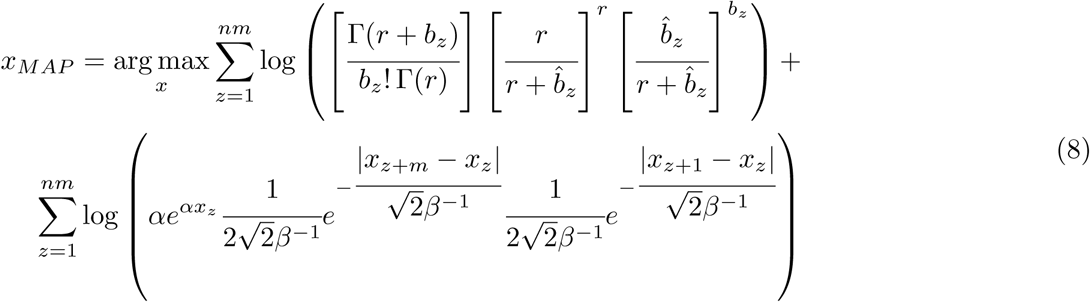

We minimize the negative of the *x*_*MAP*_ expression and ignore the hyperparameter terms since they do not depend on the pixel intensity values, *x*, and are thus constant. Further simplifying the expression and expanding the log terms we have

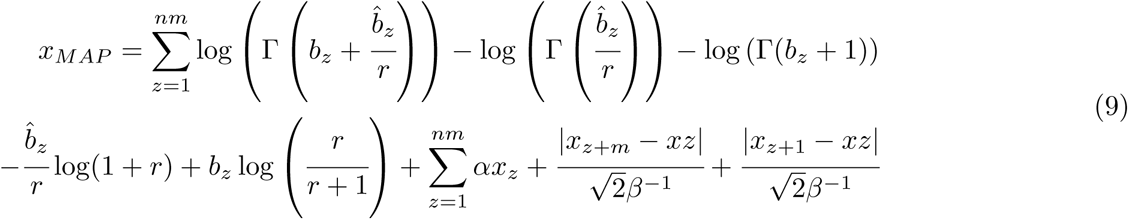

Note that we constrain the pixel intensity values to always be positive. From the above expression it is trivial to see that the exponential distribution terms and the Laplacian distribution terms simplify to the L1 norm and total variation terms respectively. This is therefore equivalent to placing a sparsity constraint on the pixel values, which intuitively makes sense as we expect most pixel values to be zero for cases of localized gene expression profiles. Total Variation minimization on the other hand has been shown to preserve edges and reduce noise in image reconstruction algorithms.

We also calculate the analytical gradient of the *x*_*MAP*_ function to speed up the convergence of the optimization algorithm. As the L1 norm and Total Variation terms are non-differentiable we use instead a differentiable variation known as the lasso (dlasso) [4]. The derivative of the *x*_*MAP*_ function is

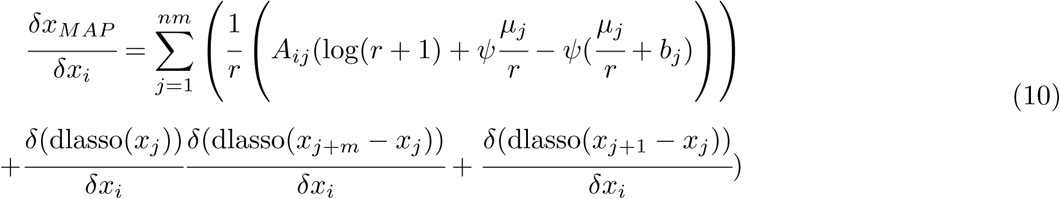

Where ψ is the digamma function

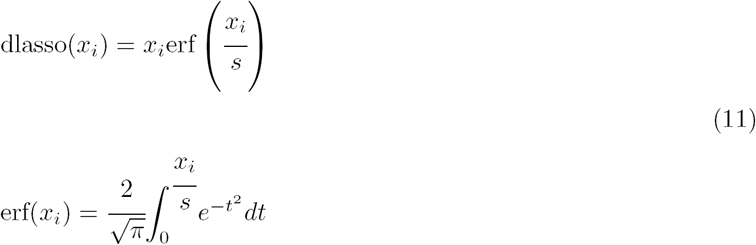

where *s* is a hyperparameter which we set equal to 0.005. The derivative of dlasso is given by:

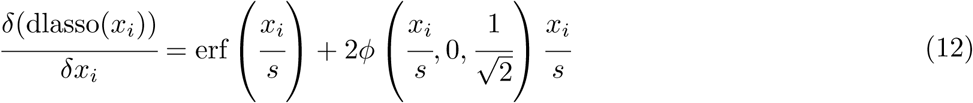

where *ϕ* is the density function of the normal distribution 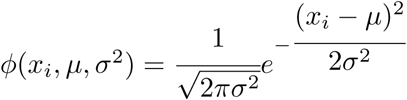

For the gradient of the total variation we follow a similar approach, calculating the difference of the neighboring pixels inside the mask.

It should be noted that 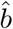 vectors are incremented by 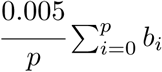 in order to prevent division by zero operations that may occur during calculation of the negative log likelihood.

#### 2.4 Computing the design matrix

To create the design matrix **A** we begin with creating a binary mask corresponding to the shape of the tissue.

We start with two micrographs (mRNA staining using FITCH-polyT) that we acquired from the sections before and after the one sequenced. We morph the two micrographs into an intermediate image by identifying corresponding features, finding a Delaunay triangulation and using it to perform a piece-wise affine transform. The same procedure was also used between the right and left section of the slice to create a symmetric image. In this way we obtain a gray-scale image which is shown as background in all the reconstructions of the paper. Finally we threshold the gray-scale image to obtain a binary mask and rescal it so that the slice width corresponds to 1 pixel.

We then use the mask to determine the position of the strips. To do this for each angle γ we consider the line bundle of slope tan(γ) and intercept *q*_*p*_. We identify *q*_1_ as the intercept of the most distant line that still overlaps with the mask. It is then possible to assign the successive *q*_*p*_ as the width of the strips is known, an stop at the last line that overlaps with the mask. Once the lines are determined *q*_*k*_ the design matrix *A*_*ij*_ can be filled accordingly.

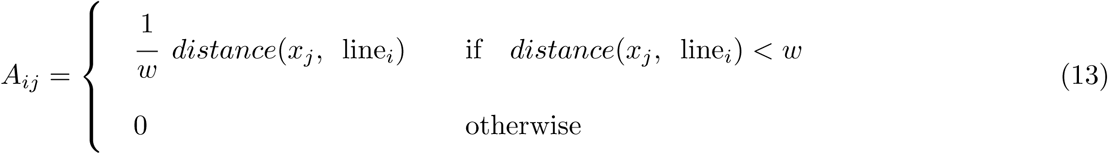

This way of defining **A** has the advantage of reducing aliasing artefacts and it is designed so that ∑_*i*_ *A*_*ij*_ = 1 *∀j* for each angle. Note that all pixels at a distance greater than *w* from the *k*^*th*^ line have contribution of zero to molecular counts of 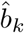.

The design matrix was modified to impose a symmetry constraint, which, in our hands, increased the accuracy of the spatial expression profiles that are recovered.

#### 2.5 K-fold cross validation and hyper-parameter optimization

The optimization problem presents two hyper-parameters: α and β. They are responsible for controlling the sparsity and smoothness of the reconstructed image, respectively. When α and β are appropriately chosen we will avoid over-fitting while also preventing relevant features from being blurred.

To score each set of hyper-parameters we adopt a k-fold cross validation procedure where all but one of the parallel-slice projections are used as training data to fit the model, and the remaining slice-projection is used as a test set to score the reconstruction by calculating the negative log-likelihood.

Note that if we had used a random subset of data points from the stacked projection the procedure we could not have avoided over-fitting as the signal in neighbouring data points and strips are correlated (i.e. training set is not independent from the test set).

A grid search is performed allowing values ranging from 0 to 10 for each hyper-parameter. In order to attain more precise values for α and β values after completion of the grid search, we use a Bayesian optimization approach (as implemented by GpyOpt library). Using values computed from the initial grid search, the Bayesian optimization procedure parsimoniously samples the cross validation function an additional number of times to determine the optimal hyper-parameter values. The hyper-parameter pair with the best log-likelihood score is selected.

### 3 Simulations and Method Evaluation

#### 3.1 Overview

In order to demonstrate the potential of the method, a number of simulated experiments were performed using ISH data from the Allen Brain Atlas. Ground truth images from the Allen Brain Atlas were up-sampled using a cubic interpolation to attain a resolution similar to reconstructions using raw data.

In all the experiments in which we added noise, we drew the noise from Poisson using the expected value of the projection **A** *·* **x** as the parameter, *λ* of the distribution, that is **b**_*sym*_ *∼* Poisson(**Ax**)

#### 3.2 Comparison to Previous Methods

Based on the Matlab code provided by the authors, we re-implemented in Python the Iterative Proportional Fitting (IPT) algorithm proposed in the Tomo-seq paper for 3D reconstruction [5]. Furthermore, to provide an example of a less under-determined problem and thus, a more fair comparison against our method, we considered a Tomo-seq adaptation that is limited to 2D sampling and reconstruction. In these experiments no noise was added to projection values.

The qualitative and quantitative differences were quantified with Pearson’s correlation coefficient, the relative 1 total error 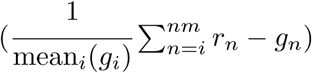 and number of pixels that differ more than a given tolerance. The last one is more precisely: 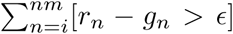 where **g** and **r** are the flattened ground truth and reconstructed images respectively and where [] are Iverson brackets. *ϵ* was chosen to be 1/16 of the difference between the maximum and minimum pixel value in the ground truth image.

#### 3.3 Influence of Angles on Reconstruction Quality

To determine the minimum number of angles to obtain a good reconstruction we considered three different levels of gene expression (average signal rescaled to 0.5, 5 and 150 counts) and presence of noise. We performed a simulation experiment on the expression pattern of the gene TAC1, reconstructions were executed using different numbers of angles. We performed the tests adding angles sequentially so that each successive simulation contained all the same cutting angles of the previous set plus one. The chosen sequence of angles was: 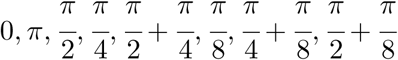. For each condition, we scored the results using the average Pearson’s R over 5 realizations.

#### 3.4 Effect of Noise at Various Expression Levels

To investigate how the quality of reconstructions varies based on gene expression, projections were scaled to various average expression levels ranging from 0.04 to 150 with additive noise sampled from a Poisson distribution. This was followed by reconstruction. The experiment was repeated for five trials for selected genes. Reconstruction quality is quantified by both Pearson’s R and PSNR.

To visualize the strength of our probabilistic algorithm, we compared the mathematical reconstructions to a reconstruction solved by a simple least-squares solver.

#### 3.5 Reconstruction Accuracy

In order to determine the accuracy of the reconstructions for different expression patterns, parallel slice-projections were simulated for 100 randomly selected genes from the Allen Brain ISH Atlas (ABA). After adding noise to the parallel slice-projection, we performed reconstructions and compared the results to the ground truth images. To recreate the conditions of the experimental data, we utilized the same angles used in the experiments for the simulations.

The comparison to ground truth images was achieved using Pearson’s R. Since the range of values corresponding to an accurate reconstruction is dependent on the dynamic range, the signal-to-noise ratio, the localization, and the sparsity of the signal, we constructed a reference distributions specific to this data.

We construct a high and low accuracy scenario by considering for each gene two gene profiles from the ABA as references: (1) for the high accuracy reference, we retrieved the most similar gene across the entire ABA and (2) for the low accuracy, we selected the gene with similarity equal to the median. More formally, being *R*_*ij*_ the pairwise correlation matrix between available genes in the Atlas, the reference distributions corresponds to the values of the vectors 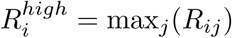 and 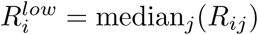.

#### 3.6 Resolution

To determine the point resolution of the technique (the distance at which the method is no longer able to discriminate two points), we used a Monte Carlo simulation approach.

We first generated a signal consisting of two points (using Gaussians PDFs) at random locations but separated at different fixed distances ranging from 0.3 to 2.0 times the strip thickness. We used the image to obtain the expected parallel slice-projections from which we draw Poisson realizations. We repeated the simulation, varying both the position of the Gaussians and the realization of the noise over the projections, and then evaluated the reconstruction results.

To determine whether the points remain distinguishable after reconstruction, two Gaussian models were fit to the images, one consisting of a single Gaussian with full covariance matrix, the second consisting of two Gaussians with diagonal covariance matrices. For each simulation, the Bayesian Information Criterion (BIC) score between the two models was computed and used as a metric of how well the two points can be discriminated.

The point resolution was then defined as the distance at which the difference in BIC scores between the individual Gaussian and mixture model becomes significantly higher than the null distribution that considers an image with only one point. In particular, the null distribution was determined with an analogous simulation but keeping the two Gaussians fully superimposed such that they form a single point. In this way, the theoretical point resolution was found to be 1.15 the strip width, with a p-value of 1.259 × 10^−5^.

### 4 STRP-seq and Data analysis

#### 4.1 Data-Preprocessing and Filtering

An important aspect of retrieving quality output reconstructions is to prepare the raw dataset through the selection of suitable genes for analysis. This section describes the two fundamental steps in the filtering process: (a) the removal of inadequate genes and (b) the selection of spatially segregated genes that contain information other than solely the total molecule count. The section goes on to describe the processing of the data immediately prior to analysis.

A total number of 24,362 and 18,580 RNA sequences with their projection profiles over five angles were provided for analysis for the mouse and lizard, respectively. It was observed that reconstruction quality tends to be poor if one of the three following conditions occurs:

- Low mean value. Genes with low expression values over projections are more affected by noise (refer to figure of simulations).
- Low non-zero counts. Genes with very low amounts of non-zero values over all projections do not allow for good estimations of the log-likelihood.
- Contradictory information between projections. Data collection may have been faulty if the quantity of non-zero values in a given angle sufficiently supersedes the quantity of non-zero values in another angle. This can pose numerical problems for the solver.

In order to filter genes based on these criteria, a threshold was required for each condition and for each data set. Genes that were below the 5^*th*^ percentile for each condition were deemed as substandard and removed from further analysis. Thresholds defined this way resulted in a minimum mean value count of 5 and a minimum non-zero count of 15 for the mouse dataset. For the lizard dataset, thresholds were a minimum mean value count of 2 and a minimum non-zero count of 10. Contradictory projections were defined to be if an angle existed that had at least 4 times fewer non-zero counts than another angle. This resulted in a new total of 5,796 and 8,183 genes in the mouse and lizard datasets, respectively. These genes are used for the comparative study described in Section 4.3.

To reduce redundancy and the amount of time to complete reconstructions, genes that had similar projection profiles to the total molecule count were removed. The projection profile of the total molecule count was processed to remove spikes via the modelling of a heteroscedastic gaussian process. The similarity between profiles was estimated by calculating the log-likelihood of each value in a given gene’s projection assuming they are samples from a negative binomial distribution where the value of the total molecule count is the expected value of the distribution. A threshold was selected based on where the scores begin to level off with respect to the distribution of their frequencies. This threshold was selected to be -1000 for the mouse data and -630 for the lizard data. This resulted in 3,880 and 2,135 genes which were then admitted for reconstruction for the mouse and lizard datasets, respectively.

Optimally chosen *α* and *β* hyperparameter values were saved for future reference. Theoretically higher *β* values correspond to flatter genes. Since the analysis performed in future sections requires spatially segregated genes, flat genes can be efficiently filtered based on the selected *β* value.

Genes that were deemed worthy of analysis were then processed in two steps: (a) application of a Gaussian blur on the image of standard deviation 2 and (b) a rescaling of pixel values to reflect the RNA molecule count from projection values for individual genes. Gaussian blurring was achieved by first determining the 50 nearest-neighbours for each pixel in a mask and multiplying their distances to the pixel in question by a Gaussian function. Values across the image were then normalized by the maximum and multiplied by the sum of RNA molecule counts across projections to reflect the gene expression intensity at a given pixel.

#### 4.2 *Mus Musculus* Analysis

##### 4.2.1 Quality of reconstructions from experimental work

In order to better define the ability of STRP-seq to mathematically reconstruct a given gene’s expression profile from raw data, Pearson’s R was calculated between the reconstruction and its ground truth profile. The ground truth profile was retrieved from the Allen Brain Atlas for genes which were present in the correct coronal layer.

The Allen Brain ISH experiments provide expression patterns in tissue which are of different dimension than that of our reference mask. In order to compare the two datasets by a correlation score, Allen brain images were rescaled and morphed using Delaunay triangulation to match the dimensions of STRP-seq gene output images. Morphing introduces an unavoidable error due to the fact that reference points for triangulation were selected manually. This implies that highly localized genes are more likely to output unreliable values for Pearson’s R, as there will be less overlap of the signal between the reconstruction and the ground truth even though the signals remain in close proximity. This can cause Pearson’s R to be negative when in fact the two spatial profiles are quite similar. In order to make the coefficient more reliable for cases such as these, a Gaussian blur was applied to the reconstructed image and original ISH image. The Gaussian blur considered the 50 nearest neighbors to determine the new value of a given pixel. Pearson’s R between ground truth and mathematically reconstructed images was then calculated using pixels strictly within the tissue mask.

##### 4.2.2 Data analysis

Given the gene expression profiles that we were able to mathematically reconstruct, we then asked if it is be possible to distinguish individual regions or cell types within the mouse brain.

In order to demonstrate this concept, three pixels were manually selected from the mouse mask such that two corresponded to regions in the thalamus and one to the cortex. Gene values were logged and then scattered against each other between thalamus-thalamus pixels and thalamus-cortex pixels. Pearson’s R was calculated for both scenarios. It should be noted that a bootstrapping approach for gene selection was used in order to dampen the effects of outliers. This resulted in an average Pearson’s R = 0.971 with standard deviation 0.0022. This method was then extended to all pixels in the mask. Using correlation coefficients which were calculated between pixels, a distance matrix was created and sorted using the Sorting Points Into Neighborhoods (SPIN) algorithm [6].

An alternative way to visualize tissue structure from gene expression profiles is through the use of a non-linear dimensionality reduction technique, such as UMAP, or a clustering algorithm. The two-dimensional UMAP projections were color-coded based on spatial location in the mask to help make sense of the distribution. The Louvain method for community detection was applied to determine whether clusters of tissue exist and support what was observed from applying SPIN to a correlation matrix between pixels.

#### 4.3 *Pogona Vitticep*s Analysis

##### 4.3.1 Independent analysis of the lizard brain

In order to select genes that were spatially localized, Principal Component Analysis (PCA) was applied to the reconstructed images to visualize basis functions that represent a significant portion of the variance of the data. Meaningful components were considered to be those that had an explained variance of at least 0.02. Since we are interested in looking for spatially varying genes, we selected genes which were not well-described by the first principal component which describes the most general distribution - the total molecule count - but correlated with other components. This filtering process resulted in the selection of 308 genes.

Relationships among pixels within the lizard tissue were investigated by calculating a pairwise correlation distance matrix and sorting it as described in Section 4.2. The individual rows in the matrix were then color-coded and mapped to the location in the brain to provide context about where the pixels are localized within the brain.

To further investigate the molecular regions of the lizard brain, the Louvain method for community detection was applied to the distance matrix. The number of neighbours and resolution parameter was set to 150 and 1.1, respectively, which permitted us to delineate four major regions in the lizard brain.

Finally, a heatmap visualizing the gene enrichment score calculated as 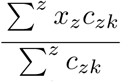 where *c*_*z*_ is a binary vector indicating if a pixel *x*_*z*_ is within the selected cluster *c*_*k*_, for individual subregions and a selection of genes was created and sorted according to hierarchical clustering. The values were then normalized over the columns. Four genes with higher enrichment scores were selected and their spatial distributions displayed.

##### 4.3.2 Comparative study of lizard and murine brain

In order to investigate the degree of similarity between the murine and lizard brains, mutual genes which were present in the datasets of both organisms were selected. This resulted in two matrices:

- The first matrix which contained murine cell types in one axis and gene values in the second axis. Cell type gene expression values were retrieved from the summary Loom file from the Zeisel Atlas ([2]; http://mousebrain.org/downloads.html) which contains expression values and metadata per cell type. Expression values were log-transformed.
- The second matrix contained gene values belonging to the lizard in one axis and pixel location in the second axis. Expression values were log-transformed.

A bootstrapping method was then performed in which 1000 genes were selected at random over 100 iterations. For each pixel in the lizard matrix, a linear regressor was fit using murine cell type gene expression levels in the mouse matrix. Regression coefficients of the cell types were stored for each pixel in the lizard mask and the median value of these 100 iterations were compared to the 10th percentile value. In cases where the median value surpasses that from the 10th percentile by at least 30%, we conclude that the regression is not considered robust and reject the result.

### 5 *Tomographer* Package

Applying the framework is facilitated by the *Tomographer* package which was developed for the mathematical reconstruction of any spatially varying signal whose input data can be provided in the specified form. The package is compatible with Python >= 3.6, and a number of external libraries are required and are listed in the Installation section of the GitHub repository at https://github.com/lamanno-epfl/tomographer. For further references, including functionality and specifications for package use, see *Tomographer* Documentation.

## Notes

https://github.com/lamanno-epfl/tomographer

### Competing Interest Statement

The authors have declared no competing interest.

